# Bidirectional synaptic plasticity rapidly modifies hippocampal representations

**DOI:** 10.1101/2020.02.04.934182

**Authors:** Aaron D. Milstein, Yiding Li, Katie C. Bittner, Christine Grienberger, Ivan Soltesz, Jeffrey C. Magee, Sandro Romani

**Author notes:** Co-corresponding authors. Please send correspondence to: Jeffrey C. Magee, Sandro Romani.

## Abstract

Learning requires neural adaptations thought to be mediated by activity-dependent synaptic plasticity. A relatively non-standard form of synaptic plasticity driven by dendritic plateau potentials has been reported to underlie place field formation in hippocampal CA1 neurons. Here we found that this behavioral timescale synaptic plasticity (BTSP) can also reshape existing place fields via bidirectional synaptic weight changes that depend on the temporal proximity of plateau potentials to pre-existing place fields. When evoked near an existing place field, plateau potentials induced less synaptic potentiation and more depression, suggesting BTSP might depend inversely on postsynaptic activation. However, manipulations of place cell membrane potential and computational modeling indicated that this anti-correlation actually results from a dependence on current synaptic weight such that weak inputs potentiate and strong inputs depress. A network model implementing this bidirectional synaptic learning rule suggested that BTSP enables population activity, rather than pairwise neuronal correlations, to drive neural adaptations to experience.

## Introduction

Activity-dependent changes in synaptic strength can flexibly alter the selectivity of neuronal firing, providing a cellular substrate for learning and memory. In the hippocampus, synaptic plasticity plays an important role in various forms of spatial and episodic learning and memory (1). The spatial firing rates of hippocampal place cells have been shown to be modified by experience and by changes in environmental context or the locations of salient features (2–8). These modifications can occur rapidly, even within a single trial (9–14). Here we investigate the synaptic plasticity mechanisms underlying such rapid changes in the spatial selectivity of hippocampal place cells.

Various forms of Hebbian synaptic plasticity have been considered for decades to be the main, or even only, synaptic plasticity mechanisms present within most brain regions of a number of species (15). The core feature of such plasticity mechanisms is that they are autonomously driven by repeated synchronous activity between synaptically connected neurons, which results in either increases or decreases in synaptic strength depending on the exact temporal coincidence (16–19). This includes so-called “three-factor” plasticity rules that, in addition to pre- and postsynaptic activity, depend on a third factor that extends the time course over which plasticity can function (15, 16, 20, 21). To implement these three-factor plasticity rules, it has been proposed that correlated pre- and post-synaptic activity drives the formation of a synaptic flag or eligibility trace (ET) that is then converted into changes in synaptic weights by the delayed third factor, usually a neuromodulatory signal (16, 22, 23).

Recently, we reported a potent, rapid form of synaptic plasticity in hippocampal CA1 pyramidal neurons that enables a *de novo* place field to be generated in a single trial following a dendritic calcium spike (also called a plateau potential) (12–14). This form of synaptic plasticity, termed behavioral timescale synaptic plasticity (BTSP), rapidly modifies synaptic inputs active within a seconds-long time window around the plateau potential. This relatively long time-course suggests that BTSP may be similar to the above-mentioned three-factor forms of plasticity, with synaptic activity generating local signals marking synapses as eligible for plasticity (ETs), and plateau potentials acting as the delayed factor that converts synaptic ETs into changes in synaptic strength. However, BTSP was shown to strengthen many synaptic inputs whose activation did not coincide with any postsynaptic spiking or even subthreshold depolarization detected at the soma (13), suggesting that changes in synaptic weight might be independent of correlated pre- and post-synaptic activity, and that BTSP may be fundamentally different than all variants of Hebbian synaptic plasticity (16-18, 24, 25). Such a non-standard plasticity rule could enable learning to be guided by delayed behavioral outcomes, rather than by short timescale associations of pre- and post-synaptic activity.

In this study, we tested the effect of dendritic plateau potentials on the spatial selectivity of CA1 neurons that already express pre-existing place fields, and therefore exhibit substantial post-synaptic depolarization and spiking prior to plasticity induction. We found that dendritic plateau potentials rapidly translocate the place field position of hippocampal place cells, both by strengthening inputs active near the plateau position, and weakening inputs active within the original place field. In order to determine if the increased post-synaptic activity in place cells is causally related to the synaptic depression observed within the initial place field, we performed a series of voltage perturbation experiments, which indicated that the direction of plasticity induced by plateau potentials is independent of post-synaptic depolarization and spiking. Next, we inferred from the data a computational model of the synaptic learning rule underlying this bidirectional form of plasticity, which suggested that it is instead the current weight of each synaptic input that controls the direction of plasticity such that weak inputs potentiate and strong inputs depress. Finally, we implemented this weight-dependent learning rule in a network model to explore the capabilities of bidirectional BTSP to adapt network level population representations to changes in the environment.

## Results

### Plateau potentials translocate existing place fields

We first examined how plasticity induced by dendritic plateau potentials changes the intracellular membrane potential (*V*_m_) dynamics in neurons already exhibiting location specific firing (i.e. place cells). Intracellular voltage recordings from CA1 pyramidal neurons were established in head-fixed mice trained to run for a water reward on a circular treadmill decorated with visual and tactile cues to distinguish spatial positions (~185 cm in length). Brief step currents (700 pA, 300 ms) were injected through the intracellular electrode for a small number (1–8) of consecutive laps to evoke plateau potentials at a second location that was between 0 and 150 cm from the initial place field (labeled “Induction 2” in Figures 1A and 1B; n=26 plasticity inductions in 24 neurons). In 8/24 neurons a “natural” pre-existing place field was expressed from the start of recording, while in 16/24 the initial place field was first experimentally-induced by the same procedure (labeled “Induction 1” in Figures 1A and 1B). In 2/24 neurons the induction procedure was repeated a third time with plateaus evoked at a different location, resulting in a total of 26 plasticity inductions in cells with pre-existing place fields.

**Figure 1.**
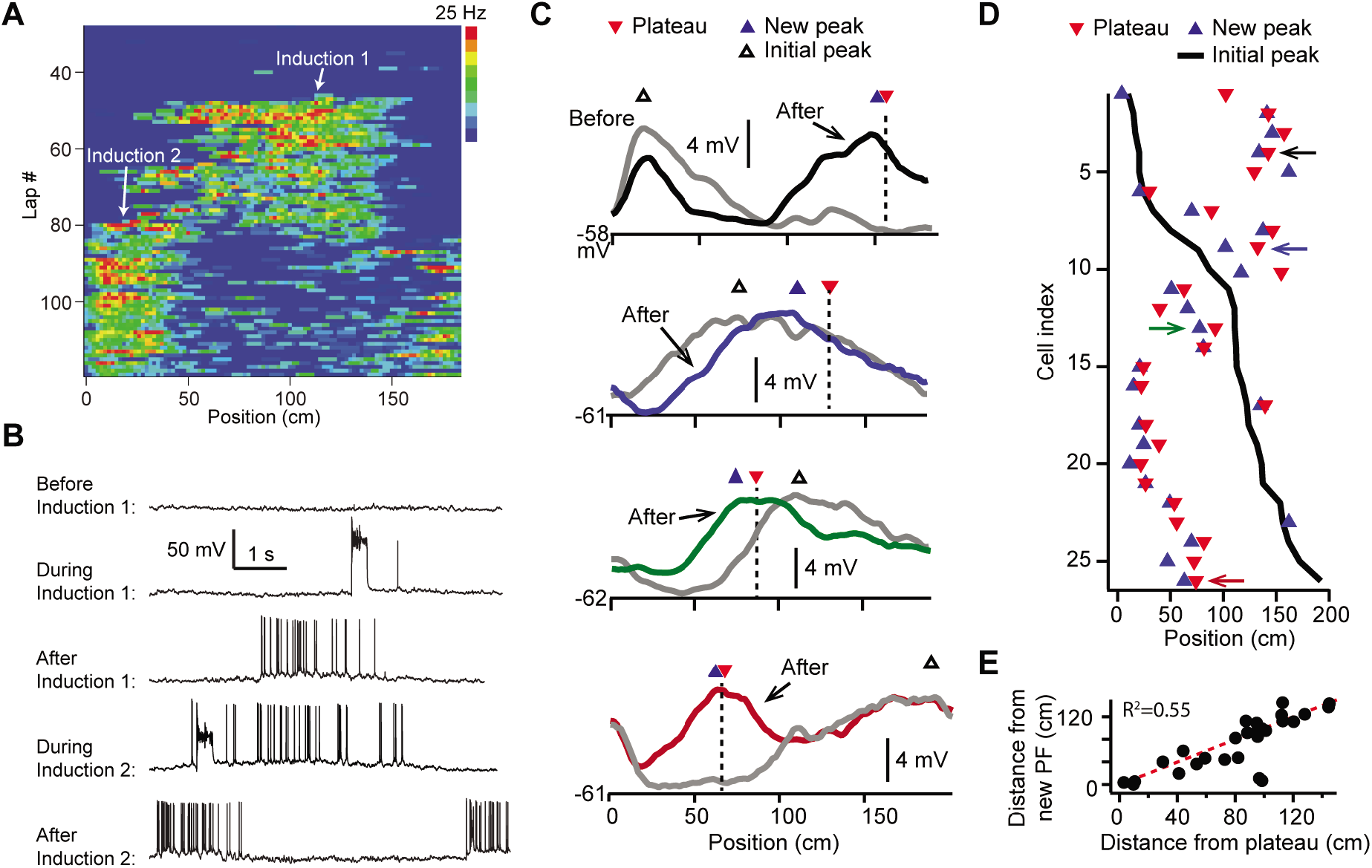
Dendritic plateau potentials translocate hippocampal place fields. (**A**) Spatial firing of a CA1 pyramidal cell recorded intracellularly from a mouse running laps on a circular treadmill. Dendritic plateau potentials evoked by intracellular current injection first induce a place field at ~120 cm (Induction 1), then induce a second place field at ~10 cm and suppress the first place field (Induction 2). (**B**) Intracellular *V*_m_ traces from individual laps in (A). (**C**) Spatially binned *V*_m_ ramp depolarizations averaged across laps before (grey) and after (black, blue, green, red) the second induction (100 spatial bins). Dashed lines and red triangles mark the average locations of evoked plateaus, black open triangles mark the location of the initial *V*_m_ ramp peak and blue triangles indicate the position of the peak of the new place field. (**D**) Data from all cells were sorted by position of initial place field. Black line indicates location of initial peak, blue triangles indicate the position of the peak of the new place field, and red triangles the position of the plateau. Neurons in (C) are indicated by like colored arrows. (**E**) The distance between the new place field and the initial place field versus the distance between the plateau and the initial place field (p=0.000015; 2-tailed null hypothesis test; explained variance (R^2^) computed by Pearson’s correlation). Red line is unity.

In most cases the evoked dendritic plateaus shifted the location of the neuron’s pre-existing place field towards the position of the second induction site (Figures 1A and 1B). Place-field firing is known to be driven by a slow, ramping depolarization of *V*_m_ from sub-to supra-threshold levels (12, 26). Isolation of these low-pass filtered *V*_m_ ramps (Figure S1A-H; Materials and Methods) revealed that plateau potentials likewise shifted the neuron’s *V*_m_ ramp towards the position of the plateau, such that the new *V*_m_ ramp peaked near the plateau position in most neurons (average distance = 19.5±4.7 cm; n=26; Figures 1C-E and S1; example cells shown in Figure 1C are indicated with matching colored arrows in Figure 1D). We also observed similar shifts in place field position to be induced by spontaneous, naturally-occurring plateau potentials in a separate set of recordings (n=5; Figures S1I-M).

### *Spatial extent of V*_m_ *plasticity*

The spatial profile of plateau-induced *V*_m_ changes (Δ*V*_m_) (Figure 2A) was obtained by subtracting the average *V*_m_ ramp for trials occurring before plateau initiation (Figure 1C; before) from the average *V*_m_ ramp for trials occurring after (Figure 1C; after). These data indicate that plateaus induced both positive and negative changes to *V*_m_ ramp amplitude (Figures 2A and 2B). In general, the increases in *V*_m_ depolarization peaked near the position of the plateau, while the negative changes peaked near the initial place field (Figures 2A, 2B, S2A and S2B). Although these changes varied considerably in magnitude across cells, the peak change in the positive direction was greater than the peak change in the negative direction (mean positive change ± SEM vs. mean negative change ± SEM: 6.73 ± 0.73 mV vs. 3.89 ± 0.32 mV, n=26 inductions; p= 0.0001, paired two-way Students t-test; Figure S2A). Aligning each Δ*V*_m_ trace to the position of the plateau (Figures 2A and 2B) demonstrates that the increases in *V*_m_ depolarization observed near the plateau position decay with distance, eventually becoming hyperpolarizing decreases in *V*_m_. At even greater distances from a plateau, Δ*V*_m_ decays back to zero (Figure 2B). To summarize the data presented thus far, dendritic plateau potentials change the location of place field firing by depolarizing *V*_m_ around the plateau position and hyperpolarizing *V*_m_ at positions within a pre-existing place field.

**Figure 2.**
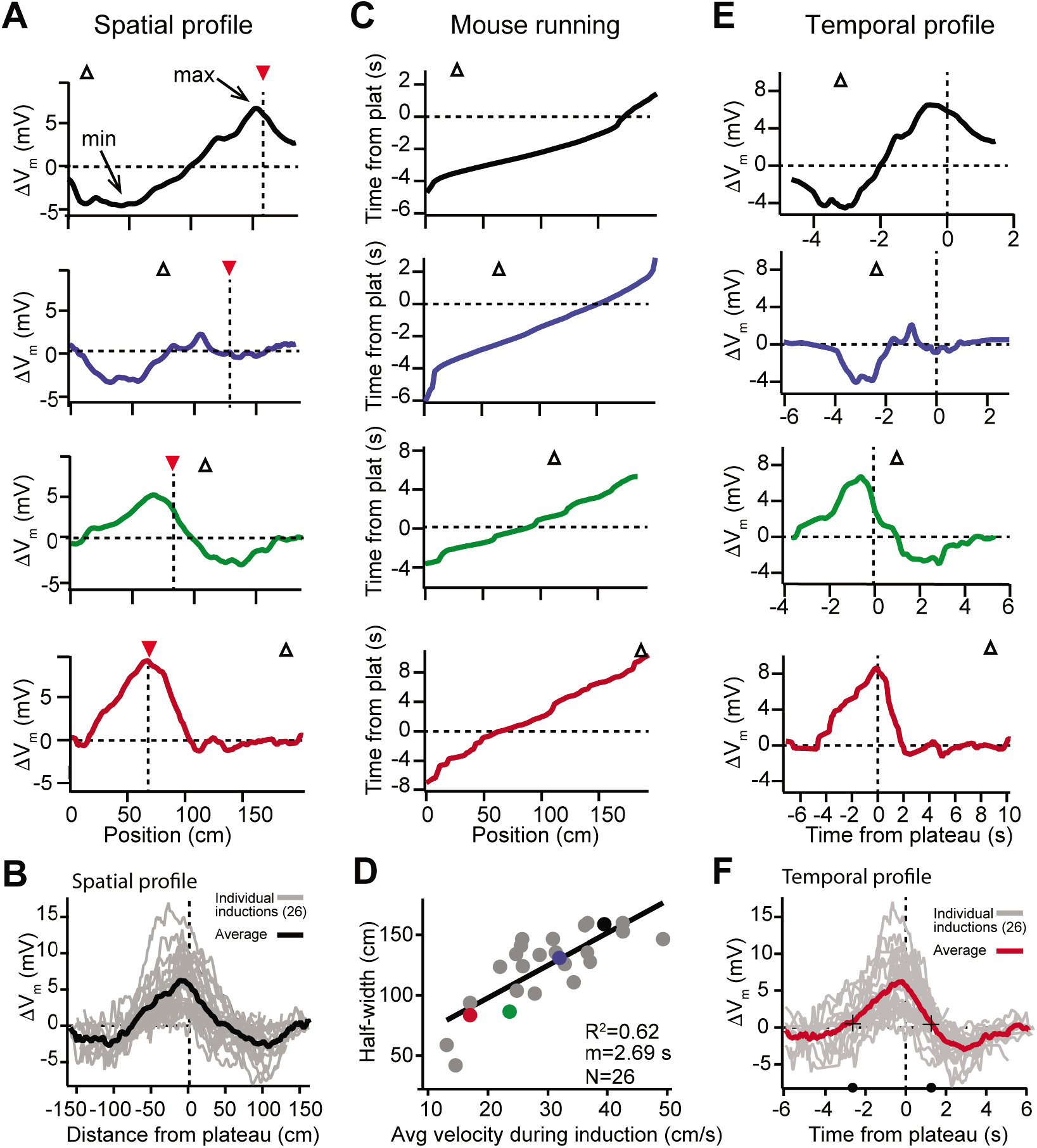
Spatial and temporal profiles of plateau-induced change in *V*_m_. (**A**) Difference between spatially binned *V*_m_ ramp depolarizations averaged across laps before the second induction and those averaged across laps after the second induction. Same example traces as shown in Fig 1C. Red triangles and dashed line indicate plateau location. Open triangles are locations of initial *V*_m_ ramp peaks. Traces have been smoothed using a 5 point boxcar average. (B) All change in *V*_m_ traces (Δ*V*_m_, not smoothed) from individual neurons (grey) and averaged across cells (black). (C) The running profile of the mice during the plateau induction trials plotted as time from plateau initiation vs spatial location (100 bins). This indicates the temporal distance of the mouse from the plateau position at any given spatial position and is used as a time base in (E) and (F). (**D**) Spatial *V*_m_ ramp half-width, calculated as distance from plateau position to the final decay of Δ*V*_m_ in a single direction, versus the average running speed of the mouse during the induction trials calculated from traces shown in (C). Individual symbols for examples shown in (A) are correspondingly colored. Grey line is linear fit (p=1.8e-06, 2-tailed null hypothesis test; explained variance (R^2^) computed by Pearson’s correlation). (**E**) Change in *V*_m_ traces (Δ*V*_m_) using the time base shown in (C). Traces have been smoothed using a 5 point boxcar average. (**F**) All change in *V*_m_ traces (Δ*V*_m_, not smoothed) from individual neurons (grey) and averaged across cells (red). Black crosses and circles indicate the 10% peak amplitude times used to calculate the asymmetry of positive changes (left/right potentiation ratio).

### *Time dependence of V*_m_ *plasticity*

Previously we showed that location-specific increases in *V*_m_ depolarization induced by plateau potentials are the result of synapse-specific increases in the strengths of spatially-tuned excitatory inputs (13). The above results suggest that, in addition to this synaptic potentiation, BTSP is also capable of inducing synaptic depression to cause location-specific decreases in *V*_m_ depolarization. In analyzing the spatial extent of the *V*_m_ changes induced by plateaus, we observed a strong linear relationship between the width of the resulting Δ*V*_m_ and the running speed of the animal during plateau induction laps (Figures 2C and 2D), which had a slope on the order of seconds. This suggested that the run trajectory of the animal (Figure 2C) affected the spatial extent of the plasticity (Figures 2A and 2B) by determining which positions were traversed within a fixed seconds-long temporal window for plasticity, as we previously reported (13). Therefore, we next analyzed the temporal relationship between plateau potentials and location-specific potentiation and depression. To do this, we used the running trajectory of the mice during plateau induction trials (Figure 2C) as a time-base for Δ*V*_m_ (Figures 2E; see also S1A-H and Materials and Methods). This analysis showed that the positive and negative changes to *V*_m_ induced in place cells occurred over a timescale of multiple seconds (Figure 2F), with the positive changes appearing to be asymmetric with respect to the onset time of the plateaus (ratio of potentiation duration before/after plateau onset: 2.2; black circles and crossmarks in Figure 2F mark the time points when Δ*V*_m_ crosses zero). This asymmetry was similar to that observed for the positive *V*_m_ changes induced by BTSP in silent cells (13). The negative changes (i.e. the hyperpolarizations indicative of synaptic depression) occurred within a time window between ±2 and ±6 seconds from the plateau in many neurons that expressed pre-existing place fields (Figures 2E and 2F). Notably, this hyperpolarization was greatly reduced, or even absent, in a set of place cells where the time delay between plateau onset and the initial place field *V*_m_ ramp was greater than 4-5 seconds (red traces in Figures 1C, 2A, and 2E; see also S2C-F), further indicating the time delimited aspect of the depression component. These data reinforce the idea that BTSP is a bidirectional form of synaptic plasticity with a seconds-long timescale that enables dendritic plateau potentials to shift the locations of hippocampal place fields by inducing both synaptic potentiation and depression.

### Plasticity drives V_m_ towards a target shape with an apparent inverse dependence on initial V_m_

We next sought to understand why dendritic plateaus induce both *V_m_* depolarization and *V_m_* hyperpolarization in cells expressing pre-existing place fields (Figures 1 and 2), but induce only *V_m_* depolarization in spatially untuned silent cells (13). Figure 3A shows that the initial temporal profile of *V_m_* in place cells with pre-existing place fields was highly variable across neurons, as plateaus were experimentally induced at different temporal distances from the existing place field in different neurons. In contrast, the change in *V_m_* (Δ*V*_m_) induced by plateaus showed a more consistent shape in time that appeared to depend on the initial level of *V_m_* depolarization at each time point prior to plasticity (Figure 3B). Large positive changes occurred at time points with relatively hyperpolarized initial *V_m_*, while time points with more depolarized initial *V_m_* were associated with less positive and more negative Δ*V*_m_. These changes resulted in final *V_m_* profiles that were highly similar across neurons, regardless of the initial *V_m_* (Figure 3C). These results indicate that BTSP induces variable changes in synaptic strength that re-shape the selectivity of neurons towards a common target shape – a place field centered near the location of evoked plateau potentials that decays towards baseline over many seconds in each direction.

**Figure 3.**
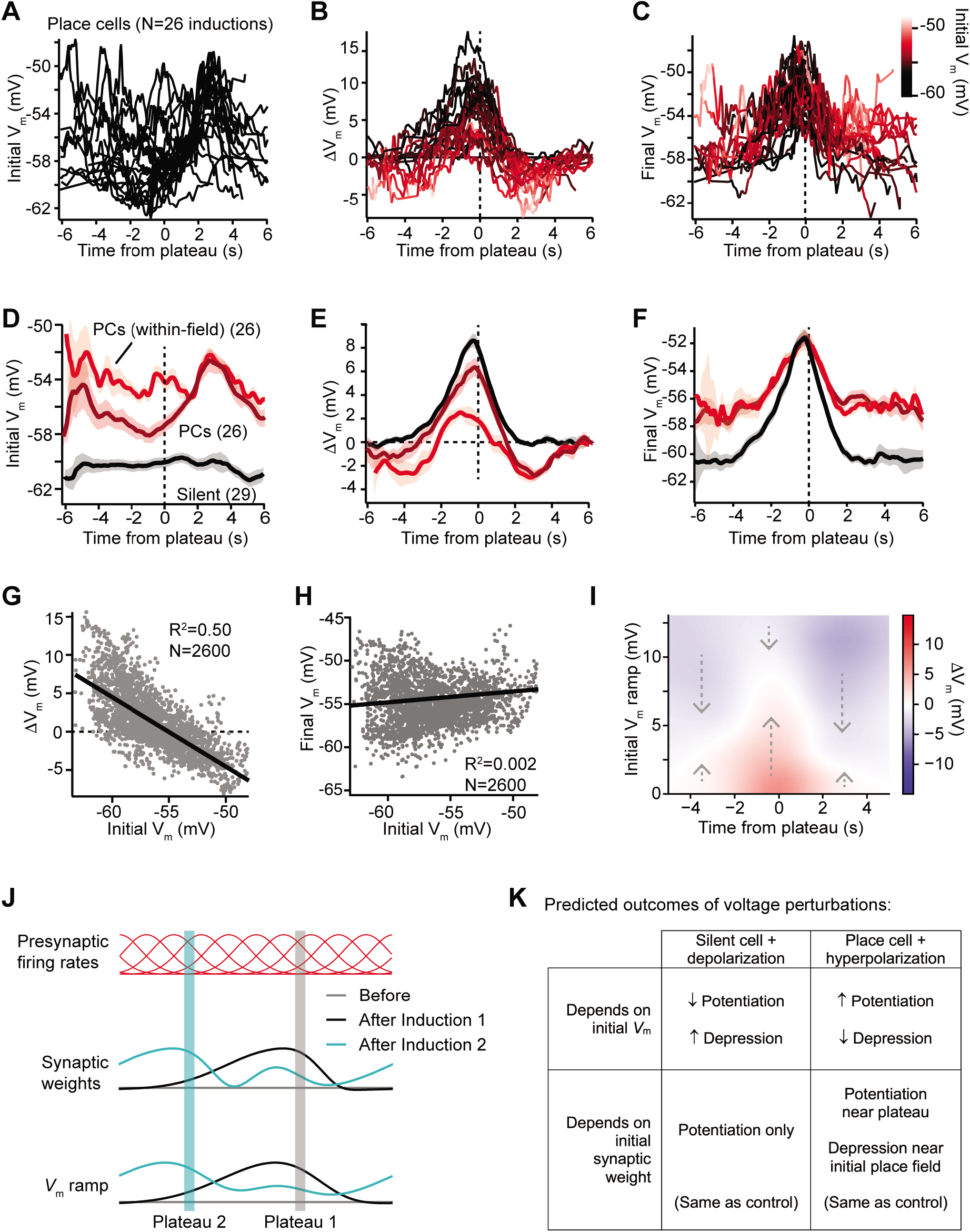
*V*_m_ ramp plasticity varies with both time delay from plateau onset and initial *V*_m_ depolarization. (**A**) Temporal profile of initial *V*_m_ before plasticity for inductions in neurons with pre-existing place fields (26 inductions from 24 place cells), aligned to the onset time of evoked plateau potentials. (**B**) Temporal profile of changes in *V*_m_ (Δ*V*_m_) induced by plasticity in all place cells. Each Δ*V*_m_ trace is color coded by initial *V*_m_. See inset color scale in (C). (**C**) Temporal profile of final *V*_m_ after plasticity in all place cells. Each *V*_m_ trace is color coded by initial *V*_m_ (color scale inset). (**D**) The temporal profiles of initial *V*_m_ before plasticity are averaged across cells and three conditions are compared: silent cells without pre-existing place fields (black), place cells (dark red), and a subset of data from each place cell at time points when each cell was more depolarized than −56 mV within its place field (light red). Shading indicates SEM across cells. (**E**) The temporal profiles of changes in *V*_m_ (Δ*V*_m_) induced by plasticity are averaged across cells and the three conditions from (D) are compared. Shading indicates SEM across cells. (**F**) The temporal profiles of final *V*_m_ after plasticity are averaged across cells and the three conditions from (D) are compared. Shading indicates SEM across cells. (**G**) Change in *V*_m_ ramp (Δ*V*_m_) plotted against initial *V*_m_ for all inductions in neurons with preexisting place fields. Black line is linear fit and correlation coefficient shown (p<0.00001, 2-tailed null hypothesis test; explained variance (R^2^) computed by Pearson’s correlation). (**H**) Final *V*_m_ ramp after plasticity plotted against initial *V*_m_ before plasticity for all inductions in neurons with pre-existing place fields. Black line is linear fit and correlation coefficient shown (p<0.014, 2-tailed null hypothesis test; explained variance (R^2^) computed by Pearson’s correlation). (**I**) Heatmap of changes in *V*_m_ ramp (Δ*V*_m_) as a function of both time and initial *V*_m_ (see Materials and Methods). Arrows indicate that the variable direction of plasticity serves to drive *V*_m_ towards the target equilibrium region (white). (**J**) Diagram depicts presynaptic spatial firing rates of a population of CA3 inputs to a postsynaptic CA1 neuron (top), the synaptic weights of those inputs before and after plasticity (middle), and the resulting postsynaptic *V*_m_ ramp, which reflects a weighted summation of the inputs. Traces are shown before (grey) and after (black) plasticity induction in a silent cell (Induction 1), and after a subsequent induction of plasticity (Induction 2, cyan) that translocates the position of the cell’s place field. (**K**) Table compares predicted outcomes of voltage perturbation experiments (depolarizing a silent cell, or hyperpolarizing a place cell), considering two possible forms of BTSP (depends on initial *V*_m_, or depends on initial synaptic weights).

We further examined this by directly comparing data from initially hyperpolarized silent cells (black in Figures 3D – F; n= 29 inductions, see Materials and Methods) to data from place cells (dark red in Figures 3D – F). We also compared these conditions to a subset of the data from place cells at time points within their initial place fields where they were relatively depolarized (greater than −56 mV) (light red in Figures 3D – F). This further supported the findings that, while changes in *V_m_* induced by plateaus were highly dependent on initial *V_m_*, these changes drove the resulting final *V_m_* ramp towards a common target shape (Figure 3F). Indeed, when all spatial bins from all place cells were analyzed, Δ*V*_m_ showed a strong inverse correlation with initial *V*_m_ (m = −0.91; Figure 3G). In contrast, final *V*_m_ showed a very weak positive correlation with initial *V*_m_ (m = 0.04; Figure 3H), which reflects that some spatial bins show no change in *V*_m_ during plasticity, either because they were traversed outside the temporal window for plasticity, or because the *V*_m_ at those positions had already reached a final *V*_m_ target value.

That BTSP induces variable changes in *V*_m_ that re-shape the *V*_m_ ramp towards a particular target shape is further evident from a heatmap depicting the relationships of Δ*V*_m_ to both initial *V*_m_ ramp depolarization and time from plateau onset (Figure 3I, positive Δ*V*_m_ in red, and negative Δ*V*_m_ in blue; see Materials and Methods). The white regions of this plot trace out a temporal profile of *V*_m_ that corresponds to the final target place field shapes shown in Figures 3C and 3F. All initial deviations from this equilibrium *V*_m_ profile resulted in either positive or negative changes to approach this target place field shape (see dashed arrows). It should also be noted that the depression of *V*_m_ in place cells appeared to be weaker than the potentiation, leaving some residual depolarization at positions distant from the peak (Figure 3F). The functional significance of this is unclear, but may suggest that BTSP induces synaptic depression at a slower rate than potentiation (27). To summarize, BTSP induces precise changes in synaptic strength that modify pre-existing place fields with any initial shape such that they approach a target shape that peaks near the location where dendritic plateaus were evoked.

### Dependence on initial Vm vs. initial synaptic weights

Altogether these data revealed that, in general, the magnitude and direction of Δ*V*_m_ depended on the time from the plateau potential, and correlated inversely with the initial *V*_m_ ramp amplitude prior to plasticity induction. Does this anti-correlation reflect a causal relationship between post-synaptic depolarization and changes in synaptic weight induced by BTSP? This possibility would require that small depolarizations induce synaptic potentiation and large depolarizations induce synaptic depression, which is actually opposite to what has been observed in CA1 pyramidal cells with a variety of other plasticity protocols (18, 28–32). Furthermore, the increased *V*_m_ depolarization within a cell’s place field also reflects the activation of strongly weighted synaptic inputs, which have been potentiated by prior plasticity (12, 13) (Figure 3J). Thus, a causal dependency on either *V*_m_ or synaptic weight could explain the data so far.

To discriminate between these two possibilities, we next devised a set of voltage perturbation experiments. We reasoned that, if increased depolarization and spiking within a cell’s place field causes synaptic depression, then artificially increasing *V*_m_ and inducing spiking in otherwise silent cells would cause plateau potentials to induce negative Δ*V*_m_. Likewise, artificially decreasing *V*_m_ and preventing spiking in place cells would prevent plateau potentials from inducing negative Δ*V*_m_ (Figure 3K). On the contrary, if the direction of plasticity depended instead on the initial strengths of synapses prior to plasticity, these voltage manipulations would have no effect on the balance between positive and negative Δ*V*_m_ (Figure 3K). It is important to note that these somatic voltage manipulations are not expected to strictly control or even completely overwhelm *V*_m_ at the synaptic sites relevant to plasticity induction (33–35) due to attenuation of current and voltage along the dendritic cable (36, 37), and compartmentalization of synaptic voltage in dendritic spines (38). However, by either increasing or decreasing the generation of somatic action potentials, this manipulation will unequivocally alter the number of action potentials that back-propagate into dendrites, which will in turn influence the activation of voltage-gated channels in dendrites and spines (e.g. Na^+^ channels, Ca^2+^ channels, and NMDA-type glutamate receptors) (33, 39). Expected changes to the mean *V*_m_ in active dendritic spines were supported by simulations of a biophysically- and morphologically-detailed CA1 place cell model expressing voltage-gated ion channels and receiving rhythmic excitation and inhibition to mimic the *in vivo* recording conditions (Figure S4) (40). Moreover, manipulation of somatic *V*_m_ and spike timing is widely used to successfully influence plasticity induction *in vitro* and *in vivo* (41–43).

According to the above scheme, we first recorded from spatially untuned silent cells, and injected current (~100 pA) through the intracellular pipette to depolarize the neurons’ *V*_m_ by ~10 mV and to increase spiking during plasticity induction trials (Figure 4A; baseline trials mean AP rate: 0.26 ± 0.25 Hz; first induction trial mean AP rate: 4.8±1.4 Hz, n=8; blue trace in Figure 4B). In all neurons tested, we observed plateau potentials to induce large positive Δ*V*_m_ at spatial positions surrounding the plateau location, and no negative Δ*V*_m_ at any spatial positions (Figure 4A; blue trace in Figure 4C). This result is inconsistent with a causal dependence on initial *V*_m_ (Figure 3K), which predicted a Δ*V*_m_ profile similar to that of control place cells at their most depolarized positions within their pre-existing place fields (red traces, “Control PCs (within-field)” in Figures 4B and 4C repeated from Figures 3D and 3E for comparison).

**Figure 4.**
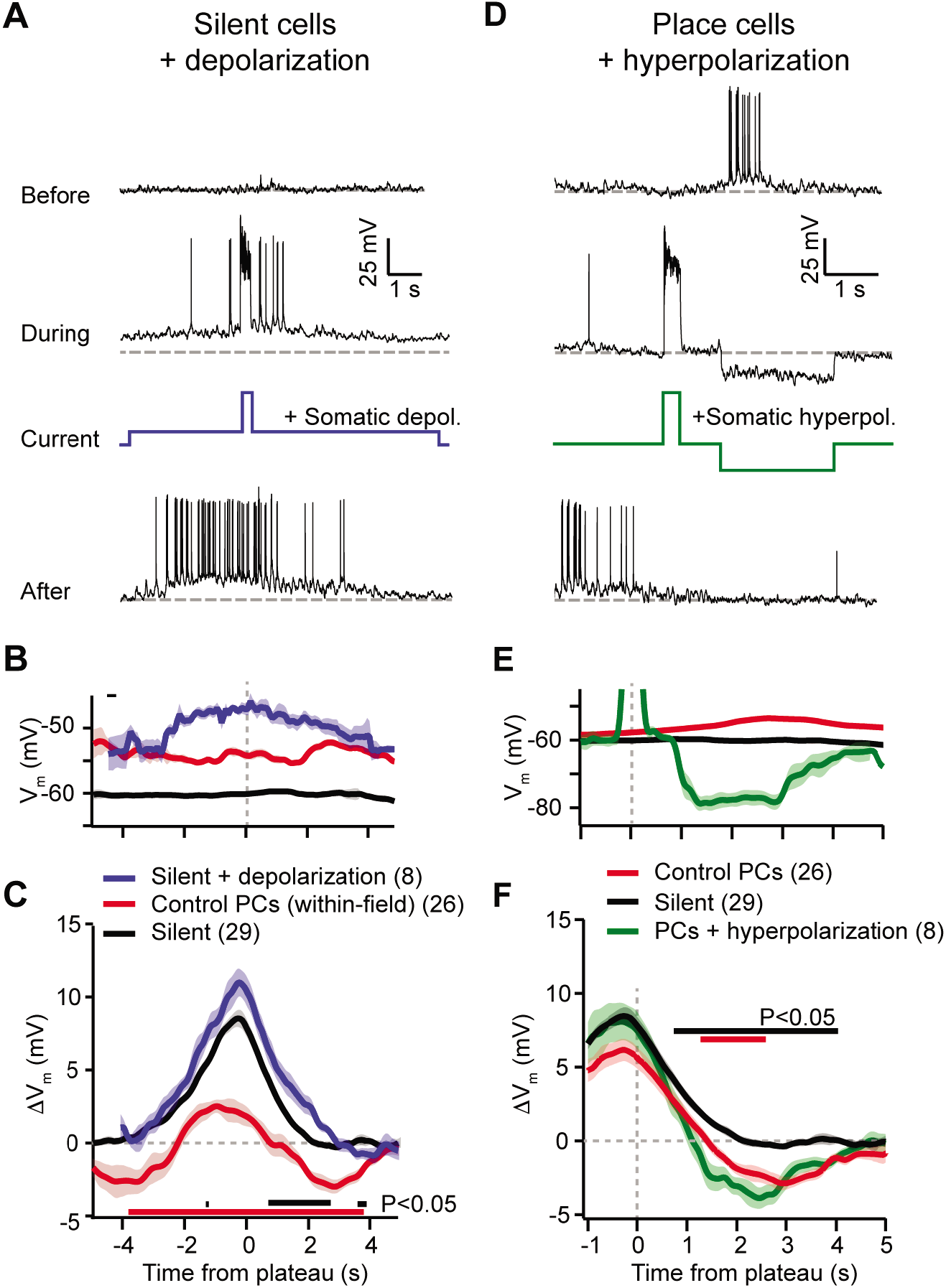
Experimental perturbation of postsynaptic activation does not change the direction of plasticity induced by BTSP. (**A**) Intracellular *V*_m_ traces from individual laps in which plasticity was induced by experimentally-evoked plateau potentials in an otherwise silent CA1 cell (top trace). During plasticity induction laps (middle trace), the neuron was experimentally depolarized by ~10 mV. Experimentally-evoked plateau potentials induced a place field (bottom). (**B**) Initial *V*_m_ before plasticity averaged across cells. Shading indicates SEM. Three conditions are compared: manipulated silent cells (Silent + depolarization; blue), data from place cells at time points within their initial place fields (Control PCs (within-field); red), and control cells without pre-existing place fields (Silent; black). (**C**) Changes in *V*_m_ ramp (Δ*V*_m_) for the same groups as in (B). Colored bars indicate statistical significance in specific time bins (p<0.05; Students two-tailed t-test). Black compares manipulated silent cells to control silent cells, and red compares manipulated silent cells to control place cells (within-field). See Materials and Methods for number of inductions in each time bin. (**D**) Intracellular *V*_m_ traces from individual laps in which plasticity was induced by experimentally-evoked plateau potentials in a place cell with a pre-existing place field (top). During plasticity induction laps, the neuron was experimentally hyperpolarized by ~25 mV at spatial positions surrounding the initial place field (middle). Experimentally-evoked plateau potentials translocated the place field (bottom). (**E**) Initial *V*_m_ before plasticity averaged across cells. Shading indicates SEM. Three conditions are compared: control place cells with pre-existing place fields (Control PCs; red), control silent cells without pre-existing place fields (Silent; black), and manipulated place cells with pre-existing place fields (PCs + hyperpolarization; green). (**F**) Changes in *V*_m_ ramp (Δ*V*_m_) for the same groups as in (E). Colored bars indicate statistical significance in specific time bins (p<0.05; Students two-tailed t-test). Black compares manipulated place cells to control silent cells, and red compares manipulated place cells to control place cells. See Materials and Methods for number of inductions in each time bin.

Next, we performed the inverse manipulation by recording from place cells and injecting current (~-150 pA) to hyperpolarize the neurons’ *V*_m_ by ~−15 mV and prevent spiking at spatial locations surrounding their pre-existing place fields while plasticity was induced at a second location (Figure 4D; baseline trials in-field mean AP rate 10.66±0.93 Hz; first induction trial infield mean AP rate 0.06±0.06 Hz, n=5; green trace in Figure 4E). This manipulation did not prevent negative Δ*V*_m_ at positions within the original place field (Figure 4D; green trace in Figure 4F), again incompatible with synaptic depression requiring elevated post-synaptic depolarization and spiking (Figure 3K). In fact, full amplitude synaptic depression was observed at locations within the original place field despite the somatic *V*_m_ being more hyperpolarized than either the silent cell (black traces in Figures 4E and 4F) or control place cell groups (red traces, “Control PCs” in Figures 4E and 4F).

These data clearly show that the direction of plasticity induced by dendritic plateau potentials is not determined by the activation state of the postsynaptic neuron. Instead, the results of these voltage perturbation experiments support the alternative hypothesis that it is the initial strength of each synapse that controls whether an input will be potentiated or depressed by BTSP (Figure 3K). However, the magnitude of potentiation and depression was slightly affected by the voltage perturbations (e.g. potentiation was slightly but significantly increased in silent cells during artificial depolarization compared to control, Figure 4C). This is consistent with the previously reported finding that BTSP induction requires activation of voltage-dependent ion channels, including NMDA-type glutamate receptors (NMDA-Rs) and voltage-gated calcium channels (VGCCs) (13), which would have predicted BTSP to depend on postsynaptic depolarization. To examine this further, we performed an additional set of experiments in which silent cells were strongly hyperpolarized by somatic current injection (~−50 mV for ~3 s just before plateau initiation) during plasticity induction (Figure S5). This manipulation decreased synaptic potentiation (Figure S5), consistent with a requirement for activation of voltage-dependent NMDARs. That such a large, non-physiological level of global *V*_m_ hyperpolarization was required to alter BTSP reinforces the finding that, operationally, the dependence is not on voltage signals associated with neuronal activation state (sustained somatodendritic *V*_m_ and action potentials), but rather on those associated with synaptic input (transient local spine depolarization) (44). Finally, these experiments do not support a role for synaptic depolarization in determining the *direction* of changes in synaptic strengths.

### Weight-dependent model of bidirectional BTSP

The above voltage perturbation experiments suggested that the form of synaptic plasticity underlying BTSP does not depend on the activation state of the postsynaptic neuron (Figure 4). This contrasts with Hebbian plasticity rules that typically depend on either the firing rate or depolarization of the postsynaptic cell to determine the amplitude and direction of changes in synaptic weight. Another difference is that BTSP appears to be inherently stable, converting synaptic potentiation into depression when input strengths exceed a particular range, whereas most models of Hebbian learning require additional homeostatic mechanisms to counteract synaptic potentiation in highly active neurons (45–49). To better understand the synaptic learning rule underlying BTSP and its functional consequences, we next sought a mathematical description of BTSP to account for the following features of the *in vivo* recording data:

1. BTSP induces bidirectional changes in synaptic weight at inputs activated up to ~6 seconds before or after a dendritic plateau potential.
2. The direction and magnitude of changes in synaptic weight depend on the initial state of each synapse such that weak inputs potentiate, and strong inputs depress.
3. BTSP modifies synaptic weights such that the temporal profile of *V*_m_ in place cells approaches a stable target shape that peaks close in time to the plateau location and decays with distance.

As mentioned previously, “three-factor” plasticity models propose a mechanism for the strengths of activated synapses to be modified after a time delay – a biochemical intermediate signal downstream of synaptic activation marks each recently activated synapse as “eligible” to undergo a plastic change in synaptic weight. This “eligibility trace” (ET) decays over a longer timescale than synaptic activation, and while it does not induce plasticity by itself, it enables plasticity to be induced upon the arrival of an additional modulatory biochemical signal. While “three-factor” models consider synaptic ETs to be generated by a coincidence of presynaptic spikes (factor 1) and postsynaptic spikes or sustained depolarization (factor 2), the results of the above voltage perturbation experiments suggest that if BTSP involves the generation of synaptic ETs, these signals depend only on a single factor – local synaptic activation. In the context of BTSP, the modulatory or “instructive signal” (IS) could be instantiated by a dendritic plateau potential. To model this, we assumed that the large magnitude dendritic depolarization associated with a plateau potential (~60 mV) effectively propagates to all synapses (50), activating an IS at each synapse and allowing a spatially and temporally local interaction between ET and IS to drive plasticity independently at each individual synapse (Figure 5A). To account for plasticity that occurs at inputs activated up to multiple seconds *after* a plateau, this IS would have to decay slowly enough to overlap in time with ETs generated after the end of the plateau (Figure 5A).

**Figure 5.**
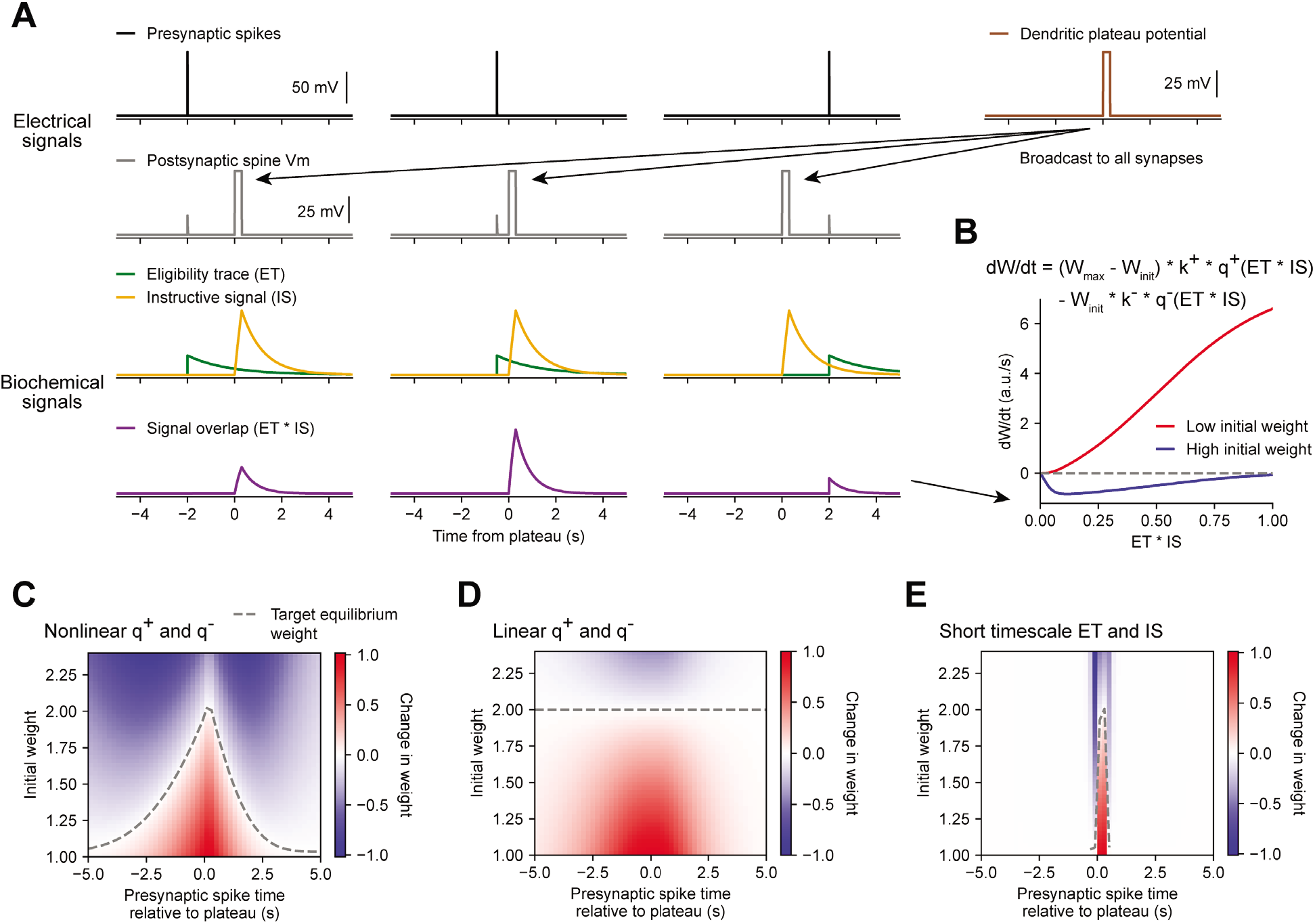
Weight-dependent model of BTSP captures essential features of plateau-induced plasticity. (**A** – **B**) Traces schematize a model of bidirectional BTSP that depends on 1) presynaptic spike timing, 2) plateau potential timing and duration, and 3) the current synaptic weight of an input before an evoked plateau. (A) Presynaptic spikes (first row, black) result in local *V*_m_ depolarization of a postsynaptic spine (second row, grey), which generates a long duration plasticity “eligibility trace” (ET) (third row, green) that marks the synapse as eligible for later synaptic potentiation or depression. The large *V*_m_ depolarization associated with the dendritic plateau potential (first row, brown) is assumed to effectively propagate to all synaptic sites (second row, grey), which generates a separate long duration “instructive signal” (IS) (third row, yellow) that is also required for plasticity. Both potentiation and depression are saturable processes that depend on the temporal overlap (product) of ET and IS (fourth row, purple). (B) Equation defines the rate of change in synaptic weight 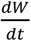 in terms of a potentiation process *q*^+^ that decreases with increasing initial weight *W_init_*, and a depression process *q*^−^ that increases with increasing *W_init_*. Plot shows the relationship between 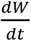 and the signal overlap *ET* * *IS* under conditions of low (red) or high (blue) initial weight. (**C** – **E**) Heat maps of changes in synaptic weight in terms of time delay between presynaptic spike and postsynaptic plateau, and initial synaptic weight for three variants of the weight-dependent model of BTSP. Dashed traces mark the equilibrium initial synaptic weight at each time delay where potentiation and depression are balanced and additional pairings of presynaptic spikes and postsynaptic plateaus result in zero further change in synaptic weight. (C) Model in which potentiation (*q*^+^) and depression (*q*^−^) processes are nonlinear (sigmoidal) functions of signal overlap (*ET* * *IS*). (D) Model in which are potentiation (*q*^+^) and depression (*q*^−^) processes are linear functions of signal overlap (*ET* * *IS*). (E) Model in which the durations of the eligibility trace (*ET*) and instructive signal (*IS*) are constrained to a short (100 ms) timescale, similar to intracellular calcium.

Accordingly, we modeled changes in synaptic weights as a function of the time-varying amplitudes of these two biochemical intermediate signals, ET and IS. For simplicity, we first considered how BTSP would change the weight *W* of a single synapse activated by a single presynaptic spike with precise timing relative to the onset of a plateau potential (Figure 5A). We modeled the synaptic ET as a signal that increases upon synaptic activation at time *t^s^* and decays exponentially with time-course *τ_ET_*:

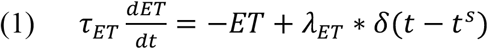

The scaling factor *λ_ET_* was chosen such that the maximum amplitude of *ET* does not exceed 1. We then modeled the IS as a signal that increases during a plateau potential with onset at time *t^p^* and duration *d* and decays exponentially with time course *τ_IS_*:

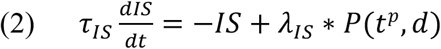

where *P* is a binary function that takes a value of 1 during a plateau and 0 otherwise. The scaling factor *λ_IS_* was chosen such that the maximum amplitude of *IS* does not exceed 1.

Next we modeled bidirectional changes in synaptic weight 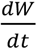 as a function of the temporal overlap or product of these two signals, *ET* * *IS*. To account for the observation that BTSP favors synaptic potentiation at weak synapses and synaptic depression at strong synapses, we expressed 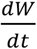 in terms of two separate plasticity processes *q*^+^ and *q*^−^ with opposite dependencies on the initial synaptic weight *W_init_*:

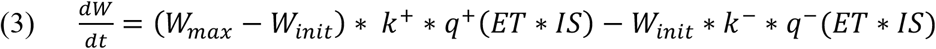

where *W* is saturable up to a maximum weight of *W_max_*, and *k*^+^ and *k*^−^ are learning rate constants that control the magnitudes of synaptic potentiation and depression per plateau potential. For simplicity, we assumed that *W* = *W_init_* throughout a trial consisting of a pairing of a single presynaptic spike with a single postsynaptic plateau potential, and we updated *W* once at the end of each trial. To calculate the net change in synaptic weight after a single trial, 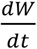 was integrated in time from the start (*t*_0_) to the end (*t*_1_) of the trial:

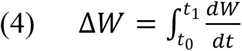

Finally, we enforced that the resulting *W* = *W_init_* + Δ*W* does not decrease below zero or increase above *W_max_*.

Experimental evidence suggests that synaptic potentiation and depression processes involve biochemical interactions between enzymes (e.g. phosphokinases like CaMKII and phosphatases like calcineurin) and synaptic protein substrates (e.g. AMPA-type glutamate receptors) (51, 52). Such concentration-limited reactions are typically saturable and nonlinear (29). Accordingly, if we define the plasticity processes *q*^+^ and *q*^−^ as saturable (sigmoidal) functions of the signal overlap *ET* * *IS* (see Materials and Methods), the resulting change in synaptic weight 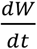 is positive and increases monotonically when initial weights are low, but is negative and nonmonotonic when initial weights are high (Figure 5B). This generates the largest negative changes in synaptic weight when signal overlap *ET* * *IS* is intermediate in amplitude. This is consistent with the *in vivo* data, which showed that negative changes in place field ramp *V*_m_ were largest at intermediate delays from a plateau (Figures 3B and 3I).

We tested this weight-dependent model of bidirectional BTSP by varying both the timing of a single presynaptic spike relative to a plateau (Figure 5A), and the initial weight of the activated synapse (Figure 5B). Model parameters were calibrated (see Materials and Methods) such that synapses with an initial weight less than a baseline weight of 1 undergo only potentiation, while synapses with higher weight undergo either potentiation or depression, depending on the timing of their activation relative to the plateau (Figure 5C). This produced a profile of changes in synaptic weight similar to the profile of changes in intracellular *V*_m_ measured *in vivo* (Figure 3I). This model also recapitulated the finding that the positive and negative changes in weight induced by BTSP appear to drive synaptic inputs towards a stable target weight, after which additional plateaus do not induce any further changes in strength (indicated in white, compare Figures 3I and 5C).

We next exploited the mathematical formulation of the model to analyze these equilibrium conditions in more detail. We defined *W_eq_* as the stable equilibrium value of W where potentiation and depression processes are exactly balanced, and the change in weight Δ*W* becomes zero:

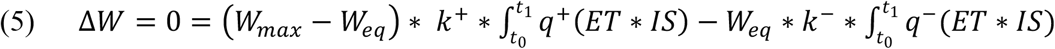

If we abbreviate the integrated potentiation and depression terms as:

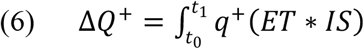

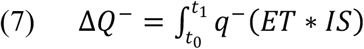

then *W_eq_* can be expressed as:

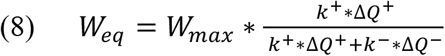

This produces a distribution of target equilibrium weights that varies depending on the timing of synaptic activation relative to a plateau (dashed line in Figure 5C), and matches the asymmetric shape of place fields induced by BTSP. In contrast, an alternative version of the model in which the potentiation and depression processes were defined to be linear instead of sigmoidal, predicted a single value for *W_eq_* regardless of the timing of synaptic activation (Figure 5D), thus failing to account for the data. Finally, we verified that the model requires long timescales for *ET* and *IS* by testing the model with shorter values (100 ms) for the decay time constants *τ_ET_* and *τ_IS_* (Figure 5E). This was unable to explain changes in synaptic weight at inputs activated at seconds-long time delays to a plateau.

Having demonstrated that this weight-dependent model of plasticity at single synapses captures the essential features of BTSP, we next tested if the model can account quantitatively for the *in vivo* place field translocation data (Figures 1 – 3). For this purpose, we assumed that the *V*_m_ ramp depolarization measured in a CA1 pyramidal cell during locomotion on the circular treadmill reflects a weighted sum of presynaptic inputs that are themselves place cells with firing rates that vary with spatial position (see Figure 3J and Materials and Methods). As a population, the place fields of these inputs uniformly tiled the track, and the firing rate of an individual input depended on the recorded run trajectory of the animal (Figure 6A, first and second rows). In this case, presynaptic activity patterns were modeled as continuous firing rates rather than discrete spike times. For each cell in the experimental dataset (n=26 inductions from 24 neurons, Figures 1 – 3), the initial weights *W_init, i_* of each presynaptic input *i* was inferred from the recorded initial *V*_m_, and the changes in weight Δ*W_μ,i_* after each lap *μ* containing an evoked plateau potential were computed as above (Equations (2) – (4)). The relevant signals modeled for an example lap from a representative cell from the dataset are shown in Figure 6A. The parameters of the model were optimized to predict the final synaptic weights (Figure 6B) and final *V*_m_ ramp (Figure 6C) after multiple plasticity induction laps (Figure S6, Materials and Methods). Across all cells, these predictions quantitatively matched the corresponding experimental data (Figure 6D). Finally, the sensitivity of changes in *V*_m_ to initial *V*_m_ and time to plateau predicted by the model recapitulated that measured from the *in vivo* intracellular recordings (Figures 6E and S7).

**Figure 6.**
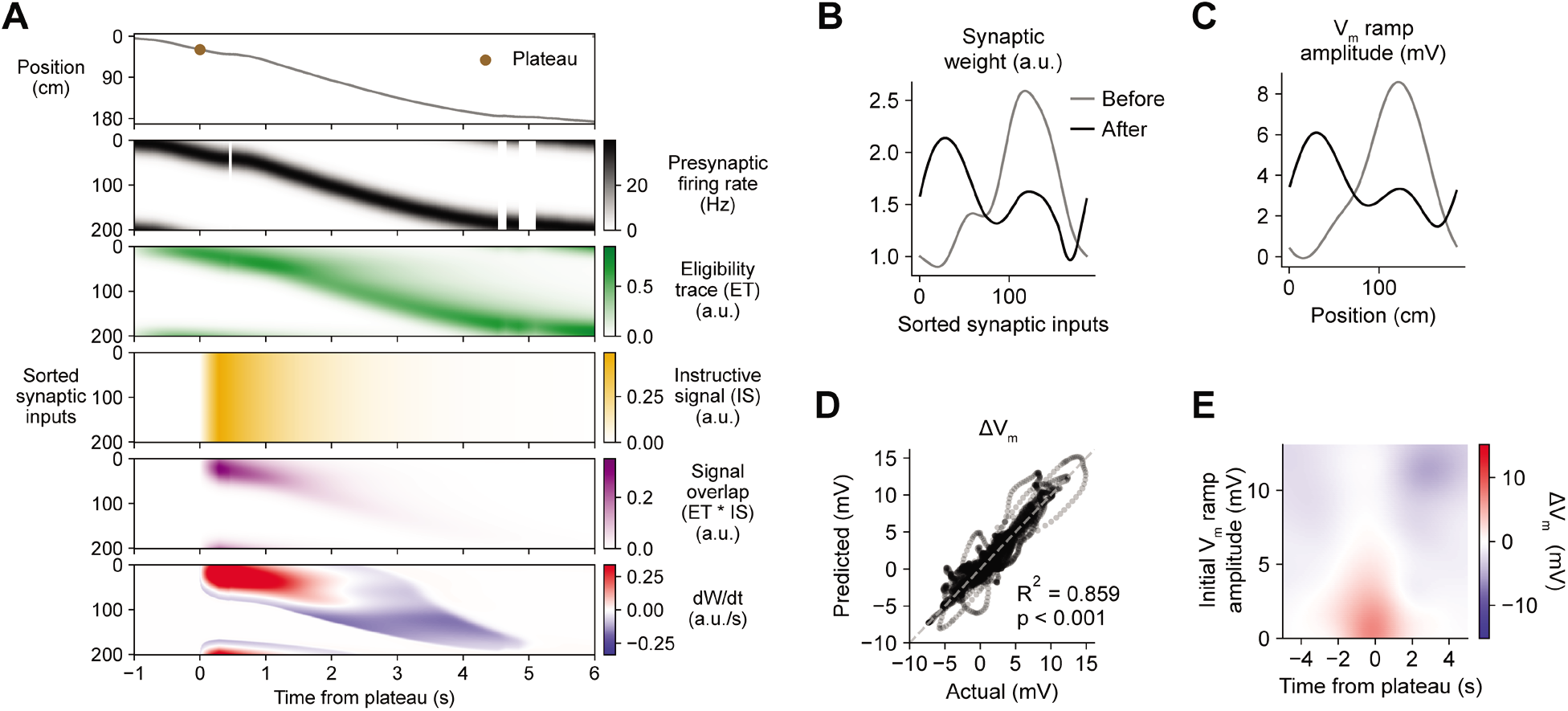
Weight-dependent model of BTSP accounts for experimentally-measured bidirectional changes in *V*_m_. (**A**) The weight-dependent model of BTSP shown in Figure 5 was used to predict plateau-induced changes in *V*_m_ in an experimentally recorded CA1 neuron given 1) the measured run trajectory of the animal during plateau induction trials (example lap shown in first row, animal position in grey), 2) the measured timing and duration of evoked plateau potentials (first row, example plateau onset marked in brown), and 3) the measured initial *V*_m_ before plasticity (shown in (C), grey). A population of 200 presynaptic CA3 place cells provided input to the model CA1 neuron. The firing rates of the presynaptic inputs were assumed to vary with spatial position and run velocity (second row, all presynaptic inputs are shown sorted by place field peak location, black). Synaptic activity at each input generated a distinct local eligibility trace (ET) (third row, green). The dendritic plateau potential generated a global instructive signal broadcast to all synapses (fourth row, yellow). The overlap between ET and IS varied at each input depending on the timing of presynaptic activity (fifth row, purple). The weight-dependent model predicted increases in synaptic weight (positive rate of change, red) at some synapses with low initial weight, and decreases in synaptic weight (negative rate of change, blue) at other synapses with high initial weight. (**B**) Synaptic weights of the 200 synaptic inputs shown in (A) before (grey) and after (black) plateau-induced plasticity. (**C**) Spatially-binned *V*_m_ ramp before (grey) and after (black) plasticity was computed as a weighted sum of the input activity. (**D**) Changes in *V*_m_ ramp amplitude (Δ*V*_m_) at each spatial bin predicted by the weight-dependent model are compared to the experimental data (n=26 inductions from 24 neurons with pre-existing place fields). Explained variance (R^2^) and statistical significance (p < 0.05) reflect Pearson’s correlation and 2-tailed null hypothesis tests. (**E**) Heatmap of changes in *V*_m_ ramp (Δ*V*_m_) predicted by the model as a function of both time and initial *V*_m_. Compare to experimental data in Figure 3I.

The above modeling results help to clarify the differences between BTSP and previously characterized forms of associative synaptic plasticity based on input-output correlations over short timescales (16, 20, 53, 54). First, the model supports the hypothesis that a dependence on initial synaptic weight is the actual source of the observed inverse relationship between initial *V*_m_ and plasticity-induced changes in *V*_m_ (Figure 3). Second, the scaling of both potentiation and depression by synaptic weight produces a balanced form of plasticity that rapidly stabilizes during repeated inductions (Figures 1, S6A and S6B) (18, 32, 46, 55, 56). Third, the time course of BTSP is determined by temporal overlap between slow eligibility signals associated with synaptic activity, and slow instructive signals associated with plateau potentials. This selects a subpopulation of synaptic inputs activated with appropriate timing to undergo a change in synaptic strength (Figure S8). Finally, instructive signals are internal signals activated by dendritic plateau potentials, rather than by spiking output, arguing that BTSP is not simply a variant of Hebbian plasticity that depends on input-output correlations over a longer timescale.

### Functional capabilities of BTSP

The above observations imply that BTSP could enable spatial representations to be shaped non-autonomously by delayed behavioral outcomes, if dendritic inputs carrying information about those outcomes are able to evoke plateau potentials (57). To evaluate the feasibility and implications of this theory, we next considered the conditions that are required for dendritic plateau potentials to be generated in the context of the hippocampal neural circuit. Previous work has shown that 1) plateau potentials are positively regulated by excitatory inputs from entorhinal cortex (12, 39, 58), 2) they are negatively regulated by dendrite-targeting inhibition (40, 58–61), 3) they occur more frequently in novel environments (62) and precede the emergence of new place fields (63), and 4) introduction of a fixed reward site induces large shifts in the place field locations of many place cells in a population, as assayed by calcium imaging (7). In order to explore the consequences of these regulatory mechanisms on memory storage by BTSP at the network level, we next constructed a network model of the CA1 microcircuit that incorporates these critical elements to regulate plateau initiation (Figure 7A), and implements the above-described weight-dependent model of BTSP (Figures 5 and 6) at each input to the network.

**Figure 7.**
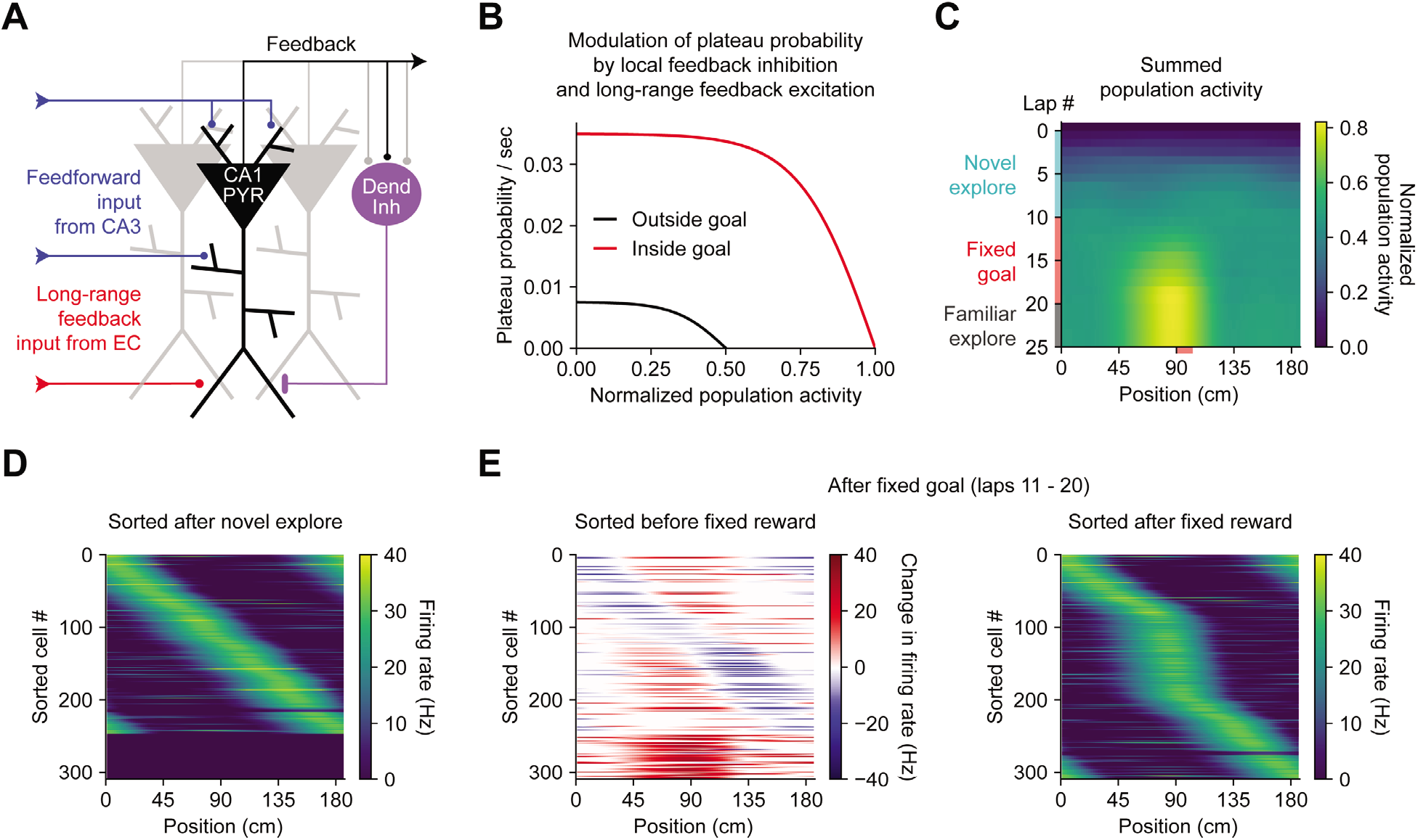
Bidirectional BTSP enables rapid adaptation of population representations in a network model. (**A**) Diagram depicts components of a hippocampal network model. A population of CA1 pyramidal neurons receives spatially tuned excitatory input from a population of CA3 place cells and a long-range feedback input from entorhinal cortex (EC) that signals the presence of a behavioral goal. The output of CA1 pyramidal neurons recruits local feedback inhibition from a population of interneurons. (**B**) The probability that model CA1 neurons emit plateau potentials and induce bidirectional plasticity is negatively modulated by feedback inhibition. As the total number of active CA1 neurons increases (labeled “normalized population activity”), feedback inhibition increases, and plateau probability decreases until a target level of population activity is reached, after which no further plasticity can be induced (black). A long-range feedback input signaling the presence of a goal increases plateau probability, resulting in a higher target level of population activity inside the goal region (red). (**C**) Each row depicts the summed activity of the population of model CA1 pyramidal neurons across spatial positions during a lap of simulated running. Laps 1-10 reflect exploration of a previously unexplored circular track. During laps 11-20, a goal is added to the environment at a fixed location (90 cm). During laps 21-25, the goal is removed for additional exploration of the now familiar environment. (**D** – **E**) Activity of individual model CA1 pyramidal neurons during simulated exploration as described in (C). (**D**) The firing rates of model neurons are sorted by the peak location of their spatial activity following 10 laps of novel exploration. A fraction of the population remains inactive and untuned. (**E**) Left: changes in firing rate of model neurons after 10 laps of goal-directed search shows place field acquisition and translocation. Right: the firing rates of model neurons are re-sorted by their new peak locations. An increased fraction of neurons express place fields near the goal position.

In a population of 500 firing-rate model CA1 pyramidal neurons, plateaus were positively regulated by a long-range feedback input from entorhinal cortex and negatively regulated by local feedback inhibition (Figures 7A and 7B) (64). Generation of plateau potentials within the population of CA1 neurons in the model was stochastic, which would result from fluctuations in inputs from entorhinal cortex that occasionally cross a threshold for the generation of a plateau potential in different cells at different times. The presence of reward delivered at a fixed goal location was implemented as an increase in input from entorhinal cortex, although an equivalent increase in plateau generation could result instead from neuromodulatory input that directly increased dendritic excitability or reduced dendritic inhibition (65–68).

During goal-directed navigation, hippocampal neurons have been shown to preferentially acquire new place fields near behaviorally-relevant locations, and to translocate existing place fields towards those locations (5–7, 69–71). We modeled this situation by simulating a virtual animal running on a circular treadmill for three separate phases of exploration (Figure 7C). At each time step (10 ms), instantaneous plateau probabilities were computed for each cell (Figure 7B), determining which neurons would initiate a dendritic plateau and undergo plasticity. During the first few laps of simulated exploration, CA1 pyramidal neurons rapidly acquired place fields that, as a population, uniformly tiled the track (Figures 7C and 7D). As neurons increased their activity over time, feedback inhibition increased proportionally and prevented further plasticity (Figures 7A–7C). During the next phase a goal was presented at a fixed location, resulting in both acquisition of new place fields nearby the goal location in a population of initially silent neurons, and translocation of place fields towards the goal location in a separate population of cells with pre-existing fields (Figures 7E, left; S9). Overall, this resulted in an increased proportion of place cells with fields near the goal position (Figure 7E, right), recapitulating experimentally observed modifications in CA1 network activity during goal-directed behavior (6). The asymmetric time course of BTSP caused the population representation of the goal in the model to peak before the goal location itself, producing a predictive memory representation of the path leading to the goal (3, 72). Simulated place cell activity remained stable in a final phase of exploration without reward (Figures 7C and S9). These network modeling results demonstrate that plasticity regulated by local network activity and long-range feedback, rather than by pairwise correlations, can enable populations of place cells to rapidly adapt their spatial representations to changes in the environment without any compromise in selectivity.

## Discussion

In summary, we observed translocation of hippocampal place fields by dendritic plateau potentials, and characterized the underlying synaptic learning rule. We found that BTSP is bidirectional, inducing both synaptic potentiation and synaptic depression in neurons expressing pre-existing place fields. The direction of plasticity is determined by the synaptic weight of each excitatory input prior to a plateau potential, and the time interval between synaptic activity and a plateau. The large magnitude of synaptic weight changes enables BTSP to rapidly reshape place field activity in a small number of trials.

The time- and synaptic weight-dependence of BTSP suggests that it is driven by an input-specific process rather than nonselective heterosynaptic (73) or homeostatic plasticity (74, 75), or modulation of cellular excitability (76, 77). A significant role for changes in inhibitory synaptic weights is unlikely given that 1) inhibitory neurons in CA1 exhibit low levels of spatial selectivity (40), 2) homosynaptic potentiation of excitatory inputs by dendritic plateau potentials can be induced with GABAergic inhibition blocked (13), and 3) inhibitory input to CA1 neurons does not change following induction of synaptic potentiation by BTSP (40).

The voltage perturbation experiments we performed (Figure 4) showed that BTSP does not depend on the activation state of the postsynaptic neuron. These results point to a fundamental difference between BTSP and existing Hebbian models of plasticity. In most previous models, including the aforementioned “three-factor” plasticity models, the firing rate (20, 53, 78, 79) or sustained level of global depolarization (30, 80, 81) at the time of presynaptic spiking primarily determines whether a synaptic weight increases or decreases (16, 47, 82). Our voltage perturbation experiments (Figures 4 and S5) show that the direction of plasticity is not determined by either global depolarization or spiking.

This lack of dependence on the postsynaptic activity or output could enable plasticity to be robust to fluctuations in postsynaptic state due to noise or network oscillations (e.g. theta or gamma) (83), and may allow the postsynaptic state to subserve other functions, such as temporal coding, without interfering with ongoing synaptic weight modifications. Furthermore, while in traditional Hebbian models of plasticity, short timescale synchrony between pre- and post-synaptic activity modifies weights to reinforce pre-existing correlations, BTSP instead provides a mechanism to either create new pairwise activity correlations “from scratch”, or remove preexisting ones based on delayed outcomes. Our network model (Figure 7) highlights how this fundamental element of BTSP could shape spatial memory storage at the network level, allowing neuronal circuits to rapidly acquire population-level representations of previously unencountered environmental features, and to modify outdated representations. This model also demonstrated that, if plateau potentials are generated by a mismatch between local circuit output and target information relayed by long-range feedback, BTSP can implement objective-based learning (84, 85).

Together our experimental and modeling results establish BTSP as a potent mechanism for rapid and reversible learning. In addition to providing insight into the fundamental mechanisms of spatial memory formation in the hippocampus, these findings suggest new directions for general theories of biological learning and the development of artificial learning systems (68, 86–89).

## Acknowledgements

We are grateful to Karel Svoboda and Wulfram Gerstner for discussions, Nicolas Brunel and Nelson Spruston for comments on the manuscript, Grace Ng for contributing to software development, Roy Phillips for behavioral device development, Ivan Raikov for technical assistance with high-performance computing, and Kristopher Bouchard at LBNL for sharing large-scale computing resources provided by the National Energy Research Scientific Computing Center, a Department of Energy Office of Science User Facility (DE-AC02-05CH11231). This work was also made possible by computing allotments from NSF (XSEDE Comet, NCSA Blue Waters, and TACC Frontera) and supported by NIH (BRAIN U19 award NS104590). SR and JM are supported by the Howard Hughes Medical Institute.

## Materials and Methods

### Animals and procedures

All experimental methods were approved by the Janelia or Baylor College of Medicine Institutional Animal Care and Use Committees (Protocol 12-84 & 15-126). All experimental procedures in this study, including animal surgeries, behavioral training, treadmill and rig configuration, and intracellular recordings, were performed identically to a previous detailed report (13) in an overlapping set of experiments, and are briefly summarized here.

*In vivo* experiments were performed in 6-12 week-old mice of either sex. Craniotomies above the dorsal hippocampus for simultaneous whole-cell patch clamp and local field potential (LFP) recordings, as well as affixation of head bar implants were performed under deep anesthesia. Following a week of recovery, animals were prepared for behavioral training with water restriction, handling by the experimenter, and addition of running wheels to their home cages.

### *In vivo* intracellular electrophysiology

Mice were trained to run on the cue-enriched linear treadmill for a dilute sucrose reward delivered through a licking port once per lap (~187 cm). A MATLAB GUI interfaced with a custom microprocessor-controlled system for position-dependent reward delivery and intracellular current injection. Animal run velocity was measured by an encoder attached to one of the wheel axles. In a subset of experiments (Figures 4 and S5), in addition to position-dependent step current to evoke plateau potentials, additional current was injected either to depolarize neurons beyond spike threshold, or to hyperpolarize neurons below spike threshold, during plasticity induction laps. While these perturbations to *V*_m_ at the soma are expected to attenuate along the path to distal dendrites (37), the pairing of back-propagating action potentials with synaptic inputs has been shown to significantly amplify dendritic depolarization (90–93). Simulations of a biophysically-detailed CA1 place cell model with realistic morphology and distributions of dendritic ion channels (40) suggest that somatic depolarization of a silent CA1 cell increases distal dendritic depolarization, and that somatic hyperpolarization of a place cell substantially reduces distal dendritic depolarization at the peak of its place field (Figure S4).

To establish whole-cell recordings from CA1 pyramidal neurons, an extracellular LFP electrode was lowered into the dorsal hippocampus using a micromanipulator until prominent theta-modulated spiking and increased ripple amplitude was detected. Then a glass intracellular recording pipette was lowered to the same depth while applying positive pressure. The intracellular solution contained (in mM): 134 K-Gluconate, 6 KCl, 10 HEPES, 4 NaCl, 0.3 MgGTP, 4 MgATP, 14 Tris-phosphocreatine, and in some recordings, 0.2% biocytin. Current-clamp recordings of intracellular membrane potential (*V*_m_) were amplified and digitized at 20 kHz, without correction for liquid junction potential. The silent-cell population of neurons (n=29) contained recordings from 17 neurons that have been previously reported (13).

### Place field analysis

To analyze subthreshold *V*_m_ ramps, action potentials were first removed from raw *V*_m_ traces and linearly interpolated, then the resulting traces were low-pass filtered (<3 Hz). For each of 100 equally sized spatial bins (~1.85 cm), *V*_m_ ramp amplitudes were averaged across periods of 5-10 minutes of running laps on the treadmill. The spatially binned ramp traces were then smoothed with a Savitzky-Golay filter with wrap-around. Ramp amplitude was quantified as the difference between the peak and the baseline (average of the 10% most hyperpolarized bins). For cells with a second place field induced, the same baseline *V*_m_ value determined from the period before the second induction was also used to quantify ramp amplitude after the second induction. Plateau duration was estimated as the duration of intracellular step current injections, or as the full width at half maximum *V*_m_ in the case of spontaneous naturally-occurring plateaus. In neurons where more than a single induction lap was present, the elapsed time between traversal of each spatial bin and onset of a plateau was potentially variable among induction laps, depending on lap-by-lap differences in run velocity. To analyze the relationship between this time interval and changes in ramp amplitude, time from plateau (e.g. Figures 2E, 2F, 3A – 3F, 3I, 4B, 4C, 4E, 4F, 6E, S1H, S1M, S2F, S3D, S5C, S5D, S7A) was conservatively estimated as the minimum time delay across all induction laps as follows: each induction lap was binned in 100 ms bins, and for each spatial position, the minimum time delay to plateau onset across all induction laps (between 1 and 8) was used to construct a single composite position versus time trace that represented the shortest induction lap in time. Since not all possible pairs of initial ramp amplitude and time delay relative to plateau onset were sampled in the experimental dataset, expected changes in ramp amplitude (e.g. Figure 3I) were predicted from the sampled experimental or model data points by a twodimensional Gaussian process regression and interpolation procedure using a rational quadratic covariance function, implemented in the open-source python package sklearn (94, 95).

*V*_m_ ramp half-width (Figures 2D and S6C) was calculated from the Δ*V*_m_ traces as the time (s) or distance (cm) between the plateau and the final return of Δ*V*_m_ to zero (or at least to 25% of min; see Figure S2G). In most cases this only occurred on one side of the plateau, during either the running period before or after the plateau. In 5/26 inductions, the mouse ran so quickly that the Δ*V*_m_ did not have time to reach 25% of min on either side of the plateau (Figure S2G), resulting in an underestimation of the ramp half-width. The average velocity was calculated from the composite induction lap (see above) as the mean velocity of the mouse from the plateau to the end of the plasticity (Figure S2).

To statistically compare Δ*V*_m_ versus time plots among groups each individual induction trace was binned in time (average of values in eighty, 100 ms, bins from −4 to +4 s). The number of points in each bin for each group are as follows: Silent cells (−4,+4s): n=19, 19, 19, 0, 20, 20, 21, 21, 25, 26, 26, 27, 27, 27, 27, 27, 28, 28, 28, 28, 29, 29, 29, 29, 29, 29, 29, 29, 29, 29, 29, 29, 29, 29, 29, 29, 29, 29, 29, 29, 29, 29, 29, 29, 29, 29, 29, 29, 29, 29, 29, 29, 29, 27, 25, 25, 25, 24, 21, 20, 19, 17, 16, 14, 14, 10, 9, 9, 8, 8, 7, 7, 7, 7, 7, 6, 6, 6, 6, 6. Silent + Depolarization (−4,+4s): n=2, 2, 4, 5, 5, 6, 6, 6, 6, 6, 6, 7, 7, 7, 8, 8, 8, 8, 8, 8, 8, 8, 8, 8, 8, 8, 8, 8, 8, 8, 8, 8, 8, 8, 8, 8, 8, 8, 8, 8, 8, 8, 8, 8, 8, 8, 8, 8, 8, 8, 8, 8, 8, 8, 8, 8, 8, 8, 8, 8, 8, 8, 8, 8, 8, 8, 8, 8, 8, 8, 8, 8, 8, 8, 8, 8, 8, 8, 8, 8, 8, 8, 8, 8, 7, 7, 6, 6, 6, 6, 5, 5, 5, 5, 3, 3. Depolarized PCs (−4 to +4s): n=6, 6, 6, 6, 6, 5, 5, 5, 4, 4, 5, 6, 6, 6, 7, 7, 6, 6, 6, 7, 7, 8, 7, 7, 7, 7, 6, 6, 5, 5, 5, 4, 4, 4, 4, 4, 4, 4, 4, 4, 4, 5, 5, 5, 5, 5, 5, 5, 6, 7, 8, 8, 8, 9, 10, 14, 14, 15, 15, 15, 15, 15, 14, 14, 14, 14, 14, 14, 14, 14, 14, 14, 14, 14, 14, 13, 12, 12, 10, 9, 9. All PCs (−1 to +4s): n=26, 26, 26, 26, 26, 26, 26, 26, 26, 26, 26, 26, 26, 26, 26, 26, 26, 26, 26, 26, 26, 26, 26, 26, 26, 26, 26, 26, 26, 26, 26, 26, 26, 26, 26, 26, 26, 26, 26, 26, 26, 26, 26, 26, 26, 26, 25, 24, 24. PCs + Hyperpolarization (−1 to +4 s): n=5, 6, 6, 7, 7, 8, 8, 8, 8, 8, 8, 8, 8, 8, 8, 8, 8, 8, 8, 8, 8, 8, 8, 8, 8, 8, 8, 8, 8, 8, 8, 8, 8, 8, 8, 8, 8, 8, 8, 8, 8, 8, 8, 8, 8, 8, 8, 8, 8, 8, 8, 8, 8, 8, 8, 8, 8, 8, 8, 8, 8, 8, 8, 8, 8, 8, 8, 8, 8, 8, 8, 8, 8, 8, 8, 8, 8, 8, 8, 8, 8, 8, 8, 8, 8. Silent + large hyperpolarization (−4 to +4): n=4, 4, 4, 4, 5, 5, 5, 6, 6, 6, 6, 6, 6, 7, 7, 7, 7, 7, 7, 7, 7, 7, 7, 7, 7, 7, 7, 7, 7, 7, 7, 7, 7, 7, 7, 7, 7, 7, 7, 7, 7, 7, 7, 7, 7, 7, 7, 7, 7, 7, 7, 7, 6, 6, 6, 6, 5, 5, 5, 5, 5, 4, 4, 4, 4, 4, 4, 3, 3, 3, 3, 2, 2, 2, 2, 2, 2, 2, 2, 2, 2.

### Quantification and statistical analysis

Statistical details of experiments can be found in the figure legends. Unless otherwise specified, measured values and ranges reflect mean ± SEM. Significance was defined as p<0.05. Sample sizes were not determined by statistical methods, but efforts were made to collect as many samples as was technically feasible. No data or subjects were excluded from any analysis.

### Computational modeling

#### Weight-dependent BTSP model

In Figures 5 and 6, we provide a mathematical model of the synaptic learning rule underlying bidirectional BTSP. In this “weight-dependent” model, the direction and magnitude of plasticity at excitatory synapses from spatially-tuned CA3 place cell inputs onto a CA1 pyramidal cell are determined by 1) the timing of presynaptic spiking relative to postsynaptic plateau potentials, and 2) the current weight of each synapse just prior to a plateau. While in Figure 5, discrete spikes were provided as presynaptic inputs to the model, in Figure 6, presynaptic inputs were provided as continuous firing rates. This model contained 9 free parameters (described in detail below), which were fit to the experimental data using an iterative, bounded, stochastic search procedure based on the simulated annealing algorithm (96, 97). This optimization sought to minimize the difference between the experimentally-recorded place cell *V*_m_ ramp depolarizations (Figures 1 – 3) and those predicted by the model (Figures 6D and 6E). Parameter optimization was considered to converge after sampling 30,000 distinct model configurations. Below we describe the model formulation in detail:

A CA1 place cell was modeled as receiving excitatory input from a population of 200 CA3 place cells with spatially tuned firing fields spaced uniformly across a ~185 cm circular track (Figure 3J). The firing rate *R_i_* of an individual input *i* with place field at position *y_i_*, depended on the recorded run trajectory of the animal *x*(*t*) (Figure 6A, first and second rows):

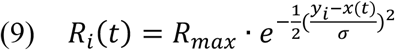

where *R_max_* is a maximum firing rate of 40 Hz at the peak of a place field, and *σ* determines the width of the place field. *σ* was set such that CA3 place field inputs had a full floor width (6 · *σ*) of 90 cm (half-width of ~34 cm) (98), though models tuned with alternative values of *σ* generated quantitatively similar predictions (“60 cm input field widths” in Figure S7). The complete run trajectory of each animal during consecutive plasticity induction laps, including pauses in running between laps, was provided as a continuous input to the model. In accordance with experimental data (12, 40), the firing rates of model place cell inputs were set to zero during periods when the animal stopped running.

The *V_m_* ramp depolarization of a CA1 place cell as a function of position, *V*(*x*), was modeled as a weighted sum of the spatial firing rates of the CA3 place cell inputs. We assumed that in silent cells prior to plasticity induction, all inputs had an initial synaptic weight of 1. This produced a background level of depolarization, *V_b_*, which was subtracted from the total weighted sum to calculate the ramp amplitude (Figures 6C – E):

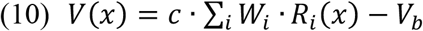

The scaling factor *c* was calibrated such that if the synaptic weights of CA3 place cell inputs varied between 1 and 2.5 as a Gaussian function of their place field locations, the postsynaptic CA1 cell would express a *V*_m_ ramp with 108 cm width and 6 mV peak amplitude, consistent with previous measurements of place field properties and the degree of synaptic potentiation by BTSP (13). For CA1 place cells already expressing a place field before plateaus were evoked at a second location, the initial synaptic weights were estimated by using least squares approximation to fit the experimentally-recorded initial *V*_m_ ramp.

At each input *i*, a postsynaptic eligibility trace *ET_i_*, was activated by presynaptic firing *R_i_*, and decayed with a seconds-long time-course (Figure 6A, third row; see also Equations (1) – (4) and text accompanying Figure 5):

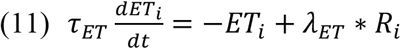

Postsynaptic dendritic plateau potentials during each induction lap *μ* with onset at time *t^p^* and duration *d* activated an instructive signal *IS* that was broadcast to all synapses and decayed exponentially with time course *τ_IS_* according to Equation (2). The duration of experimentally-induced plateaus were typically 300 ms, but spontaneous plateaus were recorded with duration up to ~800 ms.

Next, temporal overlap of eligibility traces *ET_i_*, and instructive signals *IS* were considered to drive saturable potentiation and depression processes independently at each synapse. The sensitivity of these two processes *q*^+^ and *q*^−^ to the amplitude of plasticity signal overlap was defined by generalized sigmoid functions *s*(*x, α,β*) with a scale and offset to meet the following edge constraints: *s* = 0 when *x* = 0, *s* = 1 when *x* = 1:

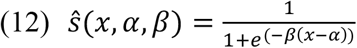

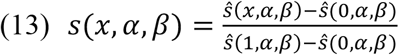

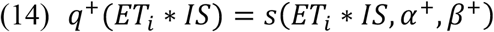

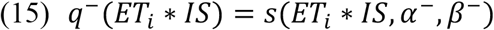

where *α*^±^ and *β*^±^ control the threshold and slope of the sigmoidal gain functions for potentiation and depression.

Finally, to capture the dependency of changes in synaptic weight 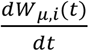 on the current value of synaptic weight *W_μ,i_* at each input *i* prior to a plasticity induction lap *μ*, we chose a two-state non-stationary kinetic model of the following form:

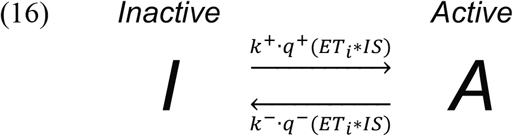

According to this formulation, independent and finite synaptic resources at each synapse occupied either an inactive state *I* or an active state *A*, and transitioned between states with rates controlled by the constants *k*^±^ and the gain functions *q*^±^ described above. The synaptic weight of each input *W_μ,i_* was defined as proportional to the occupancy of the active state *A*:

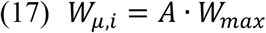

where 0 ≤ *A* ≤ 1, and *W_max_* is a free parameter controlling the maximum value of synaptic weight. Since the occupancy of each state in a kinetic model constrains the flow of finite resources between states, the net change in synaptic weight 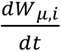 at each input *i* naturally depended on the current value of synaptic weight *W_μ,i_*:

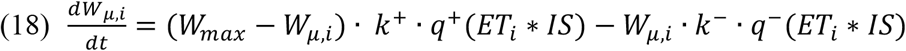

In practice, changes in synaptic weight Δ*W_μ,i_* were calculated once per lap by integrating the net rate of change of synaptic weight 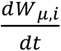 over the duration of lap *μ*. The weight-dependent model of the BTSP plasticity rule contained 9 free parameters. The range of parameter values that fit the experimental data (n=26 plasticity inductions in 24 neurons with pre-existing place fields) were as follows (mean ± SEM): 1) *τ_E_*: 863.91 ± 113.93 ms, 2) *τ_I_*: 542.76 ± 95.47 ms, 3) *α*^+^: 0.24 ± 0.05, 4) *β*^+^: 30.32 ± 6.50, 5) *α*^−^: 0.09 ± 0.04, 6) *β*^−^: 2260.61 ± 1529.97, 7) *k*^+^: 2.27 ± 0.49 /s, 8) *k*^−^: 0.33 ± 0.11 /s, 9) *W_max_*: 4.02 ± 0.17. The results of the model in response to simpler single-spike inputs in Figures 5A – 5C were obtained with the following parameter values: 1) *τ_E_*: 2500 ms, 2) *τ_I_*: 1500 ms, 3) *α*^+^: 0.5, 4) *β*^+^: 4, 5) *α*^−^: 0.01, 6) *β*^−^: 44.44, 7) *k*^+^: 1.7 /s, 8) *k*^−^: 0.204 /s, 9) *W_max_*: 5.

#### Alternative formulations of the weight-dependent BTSP model

Given the complexity of the above model, we also tested a number of alternative formulations to determine if the experimental data could be accounted for by a simpler model. First, we tested whether the filter time constants *τ_E_* and *τ_I_* that control the duration of the eligibility traces (ET) and instructive signals (IS) could be shorter by constraining their values during parameter optimization to be less than 50 ms. This model variant performed poorly in predicting the depression component of BTSP (“Short timescale ET and IS” in Figures 5E, S7A and S7B). This supports the notion that intermediate signals with durations longer than either voltage or calcium are required for the long timescale of BTSP. This also demonstrates that the nonlinear gain functions *q*^±^ are not able to compensate for shorter duration ET or IS.

Next, we determined whether the nonlinear gain functions *q*^±^ could instead be linear by replacing both the sigmoidal *q*^+^ and *q*^−^ with the identity function:

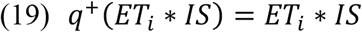

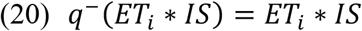

This model variant also failed to account for synaptic depression by BTSP (“Linear q^+^ and q^−^” in Figures 5D, S7A and S7B), suggesting that nonlinearity of bidirectional plasticity is an important feature of the weight-dependent BTSP model.

#### Goal-directed spatial learning model

To investigate the implications of bidirectional BTSP for reward learning by a population of CA1 place cells (Figure 7), we constructed a network model comprised of 500 CA1 pyramidal cells each receiving input from a population of 200 CA3 place cells with place fields spaced at regular intervals spanning the ~185 cm circular track. The synaptic weights at inputs from model CA3 place cells to model CA1 cells were controlled by the weight-dependent model described above (Figures 5 and 6). For this purpose, the 9 free parameters of the model were calibrated to match synthetic target *V*_m_ ramp data as follows: 1) Lap running was simulated at a constant run velocity of 25 cm/s, 2) in an initially silent cell, plasticity was induced by 3 consecutive laps with one 300 ms long plateau per lap evoked at a fixed location, 3) after plasticity, the induced place field *V*_m_ ramp had an asymmetric shape (~75 cm rise, ~35 cm decay) and a peak amplitude of 8 mV; 4) 3 additional plasticity induction laps with plateaus evoked at a location 3 seconds behind the peak location of the initial place field resulted in a 5 mV decrease in ramp amplitude at the initial peak location, and an 8 mV peak ramp amplitude at the new translocated peak position.

Before simulated exploration, all synaptic weights were initialized to a value of 1, which resulted in zero ramp depolarization in all model CA1 cells. Under these baseline conditions, each model CA1 neuron *k* had a probability *p_k_*(*t*) = *p^basal^* = 0.0075 of emitting a single dendritic plateau potential in 1 second of running. During each 10 ms time step, this instantaneous probability *p_k_*(*t*) was used to weight biased coin flips to determine which cells would emit a plateau. This stochasticity can be thought of as reflecting fluctuations in the synaptic input arriving to each cell from the long-range cortical input pathway that occasionally drive the neuron to cross a threshold for generation of a dendritic calcium spike. If a cell emitted a plateau, it persisted for a fixed duration of 300 ms, and was followed by a 500 ms refractory period during which *p_k_* (*t*) was transiently set to zero.

After the first lap, CA1 neurons that had emitted at least one plateau and had induced synaptic potentiation produced nonzero ramp depolarizations (Figure 7C). The output firing rates 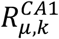 of each CA1 neuron *k* on lap *μ* were considered to be proportional to their ramp depolarizations *V_μ,k_*(*t*) after subtracting a threshold depolarization of 2 mV. The activity 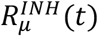 of a single inhibitory feedback element was set to be a normalized sum of the activity of the entire population of CA1 pyramidal neurons:

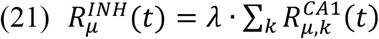

where the normalization constant *λ* was chosen such that the activity of the inhibitory feedback neuron would be 1 if every CA1 pyramidal neuron expressed a single place field and as a population their place field peak locations uniformly tiled the track. Then, the probability that any CA1 neuron *k* would emit a plateau *p_k_* (*t*) was negatively regulated by the inhibitory feedback term 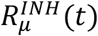:

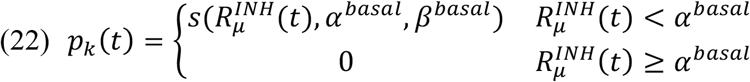

where *α^basal^* defined a target normalized population activity (set to 0.5) and *β^basal^* defined the slope of a descending sigmoid function with a maximum value of 0.0075 (Figure 7B).

In some laps, a specific location was assigned as the target of a goal-directed search. To mimic an increase in the activity of the long-range input from entorhinal cortex signaling the presence of the goal, the probability that a CA1 neuron would emit a plateau potential *p_k_*(*t*) was transiently increased when the simulated animal crossed the goal location for a period of 500 ms. Within the goal region, the relationship between *p_k_*(*t*) and 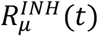 was instead:

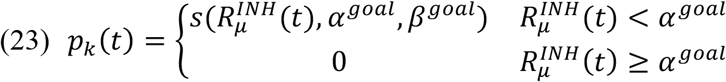

where *α^goal^* is an elevated target normalized population activity (set to 1.0) and *β^goal^* defines the slope of a descending sigmoid function with a maximum value of 0.035, corresponding to an elevated peak plateau probability (Figure 7B).

### Data and code availability

The complete dataset and Python code for data analysis and model simulation is available at https://github.com/neurosutras/BTSP. Additional MATLAB and Igor acquisition and analysis scripts will be made available upon request.

## Supplementary Figures

**Figure S1.**
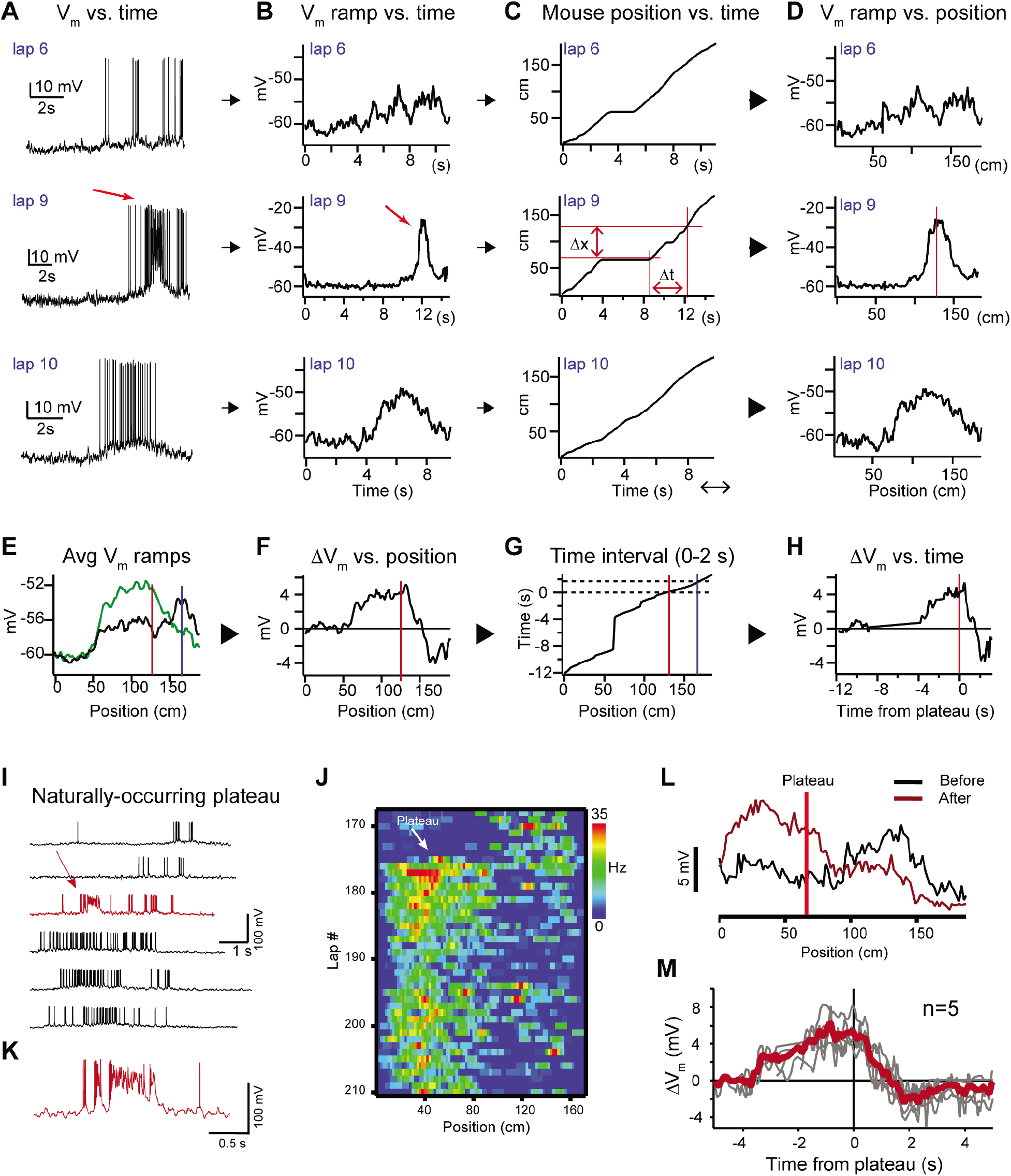
Characterization of BTSP-induced changes in *V*_m_ (related to Figure 1). (**A**) Raw *V*_m_ traces of three laps (6, 9 & 10) from a neuron that expressed a naturally-occurring place field (on laps 1-8), a naturally-occurring plateau potential (on lap 9), and a new place field on the subsequent laps (laps 10-35). (**B** – **D**) The action potentials are removed and the resulting traces are smoothed to generate *V*_m_ ramps (B) (see Materials and Methods). The position of the mouse in time (C) is used to convert the *V*_m_ ramps from time to position (D). Red lines in (D) indicate position of plateau. Red lines in (C) indicate temporal (Δt) and spatial (Δx) distance from the plateau location to an example position. (**E**) Average spatially-binned *V*_m_ ramp for trials before (laps 1-8, black) and after the plateau (laps 10-19, green) (100 spatial bins). Red line indicates plateau position, blue line indicates peak location of initial existing place field. (**F** – **H**) The difference between the before and after *V*_m_ ramps (F) is converted back to time (H) using the position of the animal in time during the plateau trial (G). Dashed black lines in (G) indicate time interval between the plateau position and the position of the initial place field. (**I**) Example *V*_m_ traces for laps before and after (black) a naturally-occurring plateau potential (red) in a CA1 cell with a pre-existing place field. (**J**) Spatial firing rate heat map shows naturally-occurring plateau shifted pre-existing place field. (**K**) Expanded view of plateau in (I). (**L**) Spatially binned subthreshold *V*_m_ ramp depolarizations averaged across laps before (black) and after (dark red) a naturally-occurring plateau (light red line). (**M**) Changes in *V*_m_ ramp versus time to plateau for place cells in which a single naturally-occurring plateau reshaped a pre-existing place field (n = 5). Naturally-occurring plateaus induced both potentiation and depression that resulted in translocation of the place field.

**Figure S2.**
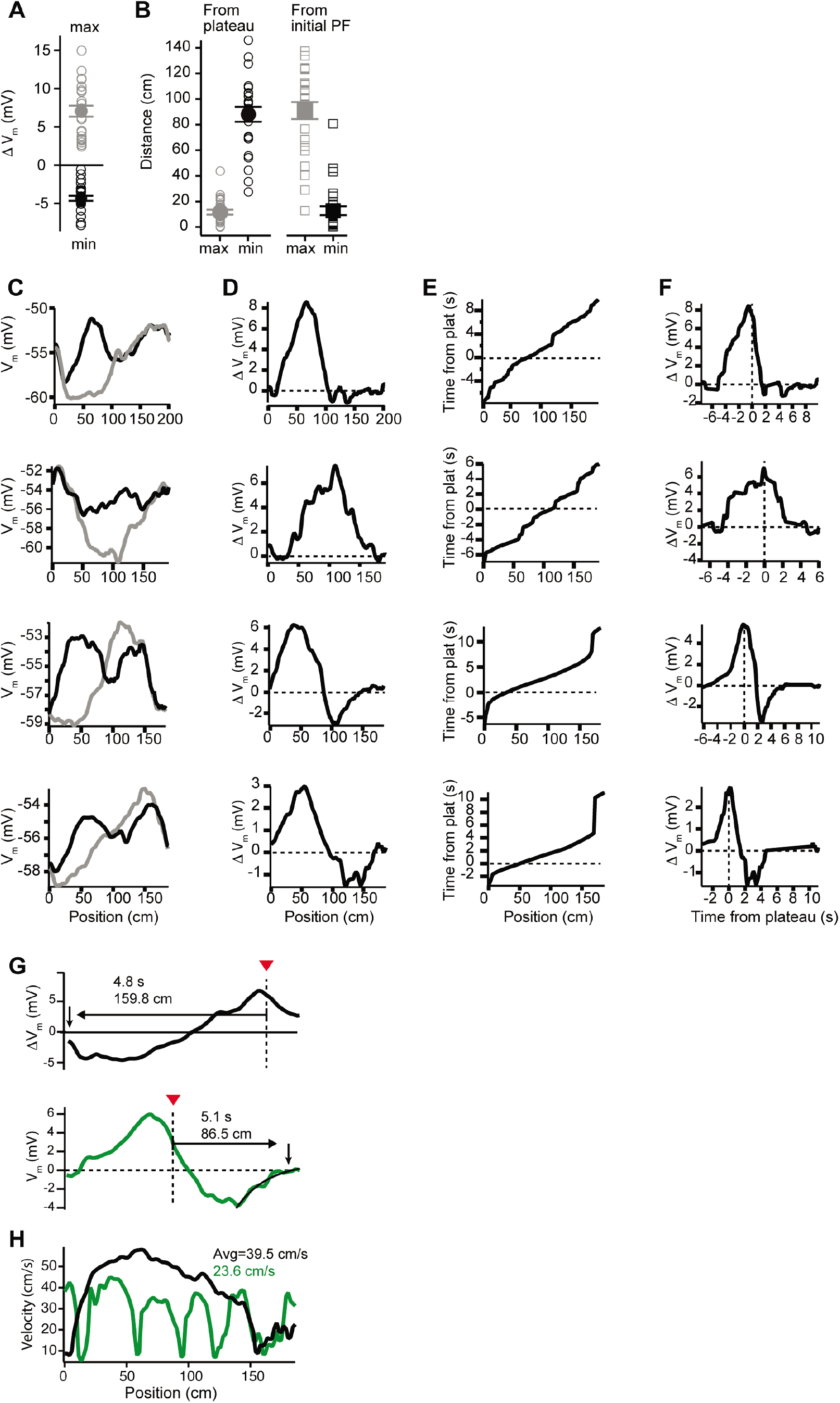
*V*_m_ changes in space and time in place cells with pre-existing place fields (related to Figure 2). (**A**) Peak positive changes in *V*_m_ ramp (max, grey open circles) and peak negative changes (min, black open circles) for all cells. Mean and SEM are indicated with filled circles and bars. (**B**) Left: the distance from the plateau for the position with peak positive change (max, grey open circles) and for the position with peak negative change (min, black open circles). Mean and SEM are indicated with filled circles and bars. Right: The distance from the peak location of the initial place field position for the position with peak positive change (max, grey open circles) and for the position with peak negative change (min, black open circles). Mean and SEM are indicated with filled circles and bars. (**C**) *V*_m_ ramp before (grey) and after (black) plateau for neurons with large temporal separation between plateau and initial place field locations. (**D**) Change in *V*_m_ induced by plateau. (**E**) The running profile of the mice during the plateau induction trials plotted as time from plateau initiation vs spatial location. (**F**) Change in *V*_m_ induced by plateau plotted using time-base calculated from induction trials. (**G**) Changes in *V*_m_ ramp for two example cells from Figures 1 and 2. Distance and time from plateau (dashed line) to the end of plasticity (down pointing arrow) are shown. Note upper black trace does not return to zero before lap is complete. Lower green trace also shows the exponential fit of decay used to determine zero crossing location. (**H**) Animal run velocity during induction trials is shown for the two example cells in (G). Average velocity values between plateau location and the zero-crossing location are indicated for each cell.

**Figure S3.**
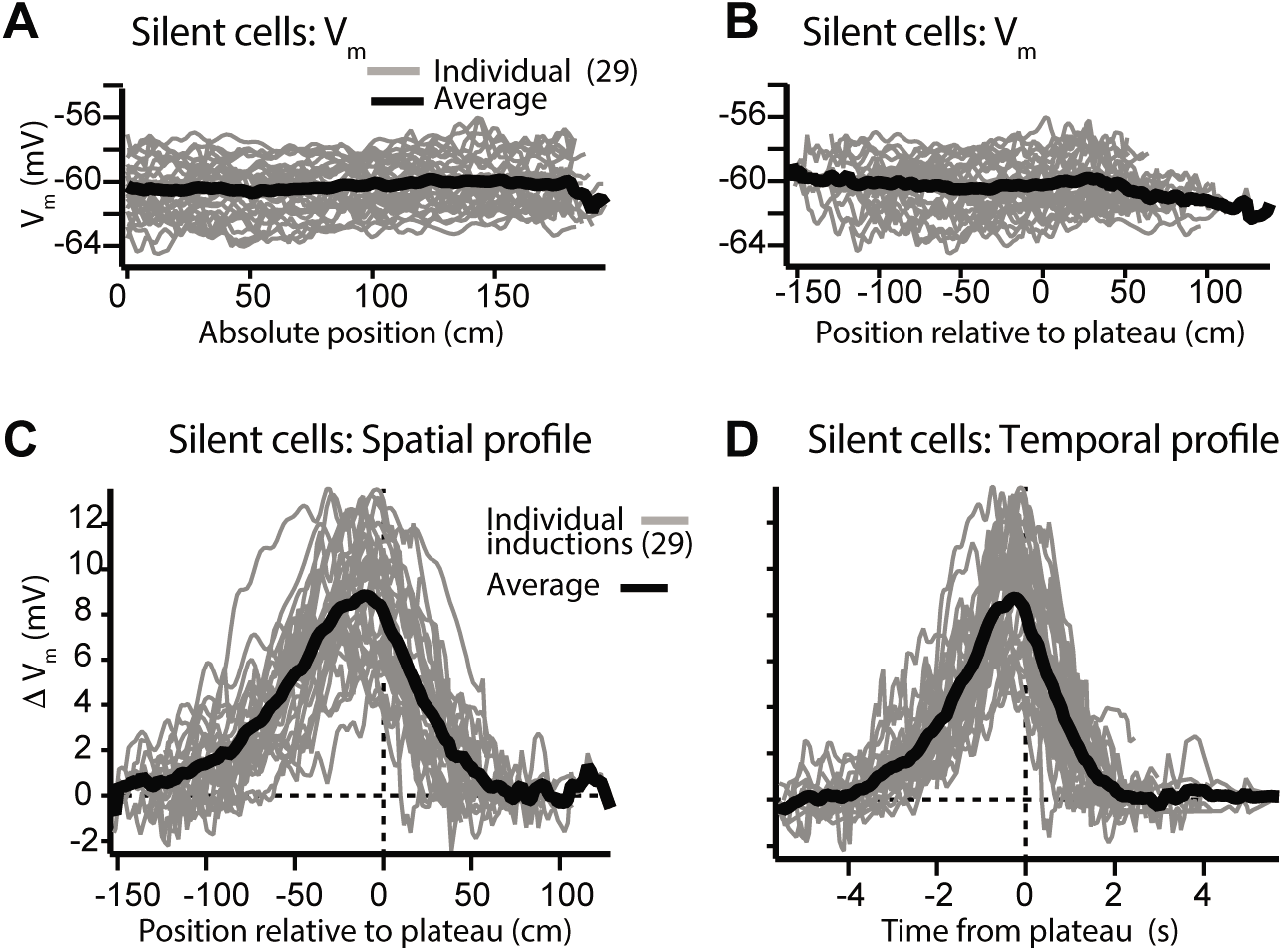
*V*_m_ changes in space and time in silent cells without pre-existing place fields (related to Figure 3). (**A**) Spatially-binned initial *V*_m_ traces for silent cells that did not express pre-existing place fields (individual traces, grey; average across cells, black) (100 spatial bins). (**B**) Same as (A) but aligned to location of plateau. (**C**) Spatial profile of change in *V*_m_ (Δ*V*_m_) for silent cells. (**D**) Temporal profile of Δ*V*_m_ for silent cells.

**Figure S4.**
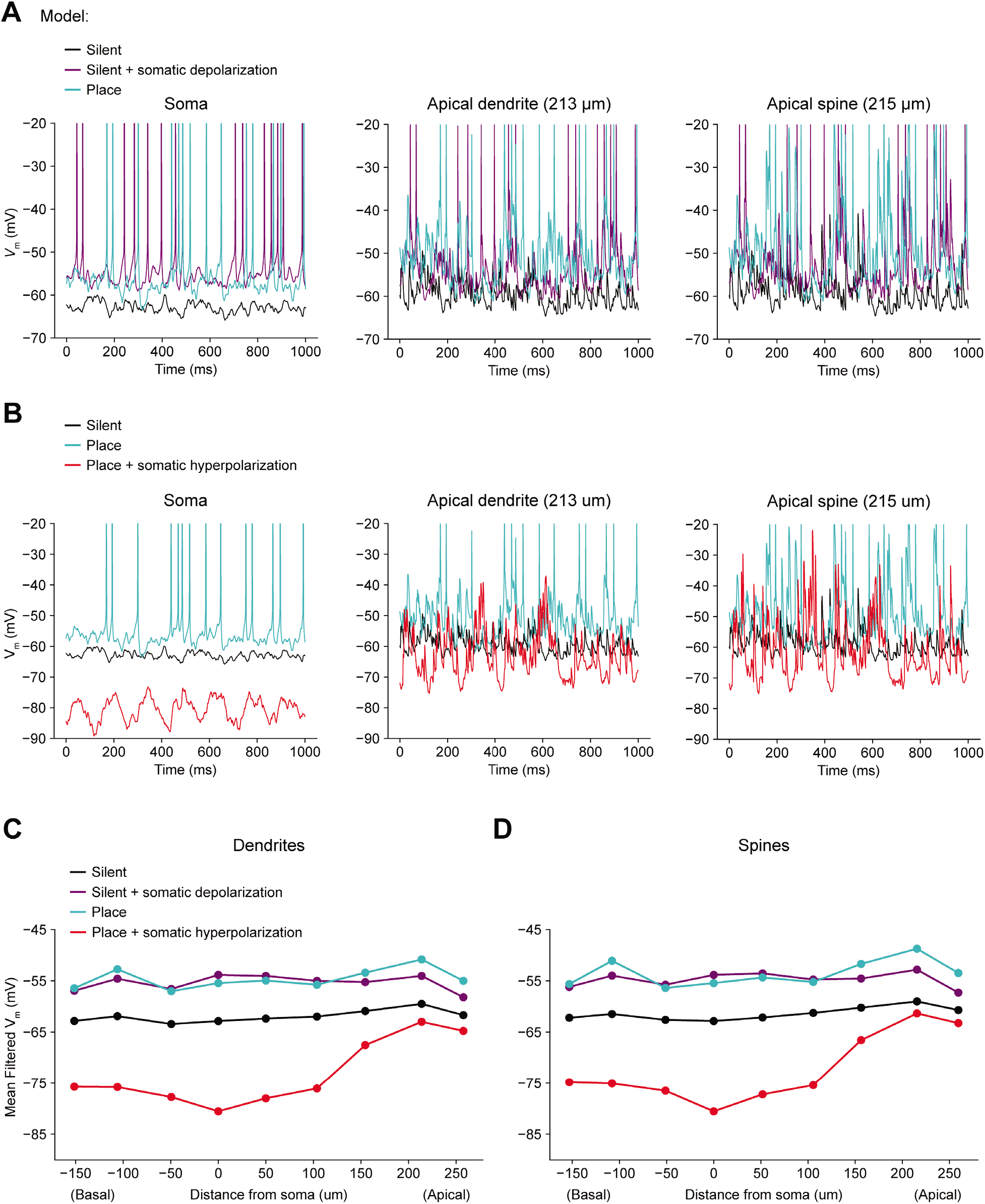
Biophysically-detailed simulations of depolarizing and hyperpolarizing somatic *V*_m_ perturbation experiments (related to Figure 4). (**A**) Simulation of a biophysically-detailed CA1 pyramidal cell model with realistic morphology and dendritic ion channel distributions (Grienberger et al., 2017) to estimate the effect of somatic depolarization on distal dendritic *V*_m_ (c.f. Figure 4). Three conditions are compared: a silent cell with uniform input weights (black), a silent cell with ~10 mV of depolarization induced by somatic current injection (purple), and a place cell receiving potentiated inputs at the peak of its place field (blue). Simulated *V*_m_ traces were recorded from soma (left), distal apical oblique dendrite (center), and a distal apical dendritic spine (right). Under these conditions, a combination of attenuated depolarization and back-propagating action potentials invades dendrites and amplifies local synaptic input by activating dendritic voltage-gated ion channels, resulting in a level of dendritic depolarization comparable to the place field condition. (**B**) Simulations estimate the effect of somatic hyperpolarization on distal dendritic *V*_m_. Three conditions are compared: a silent cell with uniform input weights (black), a place cell receiving potentiated inputs at the peak of its place field (blue), a place cell with ~25 mV of hyperpolarization induced by somatic current injection (red). Under these conditions, action potentials are prevented, and attenuated hyperpolarization invades dendrites, but local synaptic input continues to drive substantial intermittent local depolarization. (**C**) Mean low-pass filtered *V*_m_ at simulated dendritic recording sites at varying distances from the soma. (**D**) Same as (B) for simulated recordings from dendritic spines.

**Figure S5.**
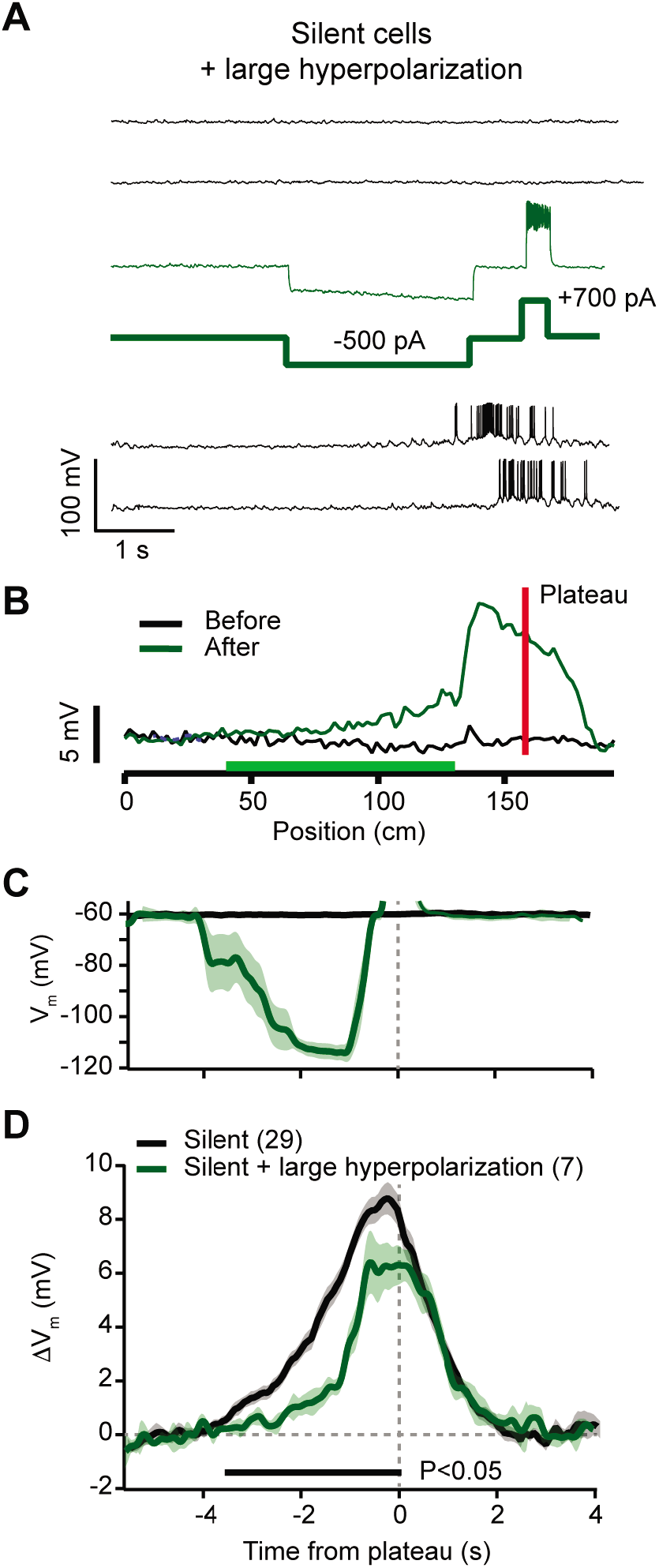
Hyperpolarization of silent cells during plasticity induction reduces synaptic potentiation (related to Figure 4). (**A**) Top black traces are *V*_m_ during laps before induction in a silent cell. During plasticity induction (thin green trace) somatic current injection (−500 pA, thick green trace) hyperpolarized *V*_m_ by ~40 mV for a duration of ~3 s ending 500 ms before the plateau induction location (+700 pA, thick green trace). Bottom black traces are *V*_m_ during laps after plasticity induction. (**B**) Spatially binned subthreshold *V*_m_ ramp depolarizations averaged across laps before (black) and after (green) plasticity induction. Red line indicates plateau location, green bar indicates locations of hyperpolarizing current injection. Note relatively small potentiation at locations with hyperpolarizing current. (**C**) Average *V*_m_ for control silent cells without pre-existing place fields (Silent; black), and the manipulated neurons (Silent + large hyperpolarization; green). (**D**) Average change in *V*_m_ ramp amplitude for the same groups as in (C). Black bar indicates time bins with statistically significant difference between manipulated and control groups (p<0.05; Students two-tailed t-test). See Materials and Methods for number of inductions in each time bin.

**Figure S6.**
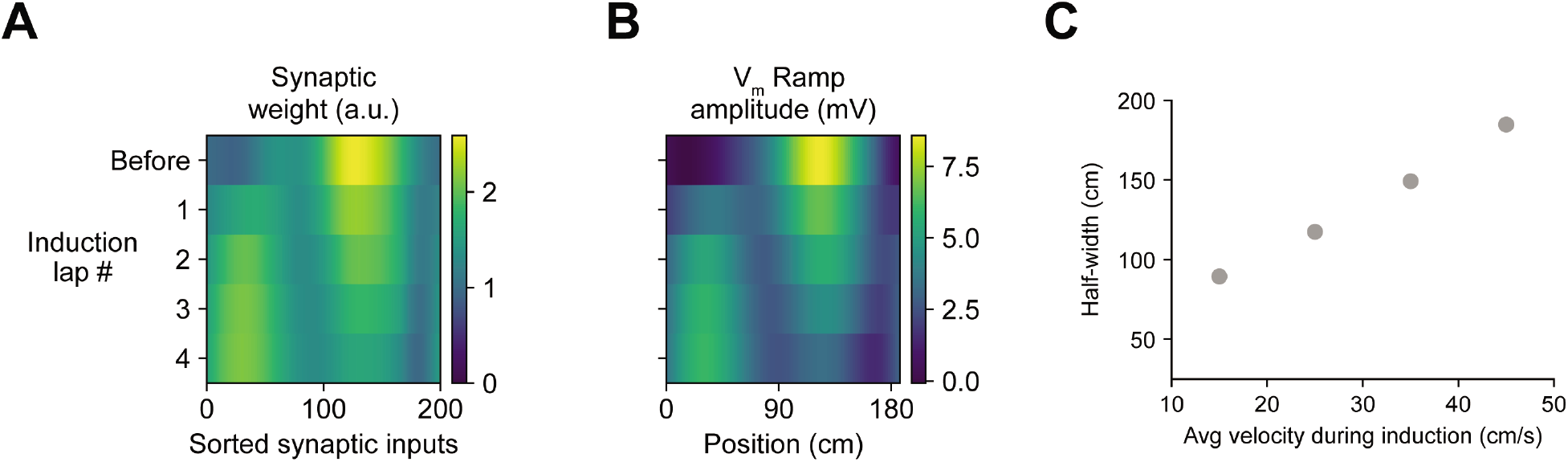
Sensitivity of induced place field *V*_m_ ramps to repeated plateau potentials and run velocity in the weight-dependent model of BTSP (related to Figures 2 and 6). (**A**–**B**) For the example cell shown in Figures 6A – C to illustrate the weight-dependent model of BTSP described in Figure 5, the values of synaptic weights (A) and *V*_m_ ramp amplitude (B) before and after each of four plasticity induction laps are shown. (**C**) Simulations of plasticity induction under conditions of constant run velocity reveals a positive relationship between run velocity and spatial width of the induced place field (compare to Figure 2D).

**Figure S7.**
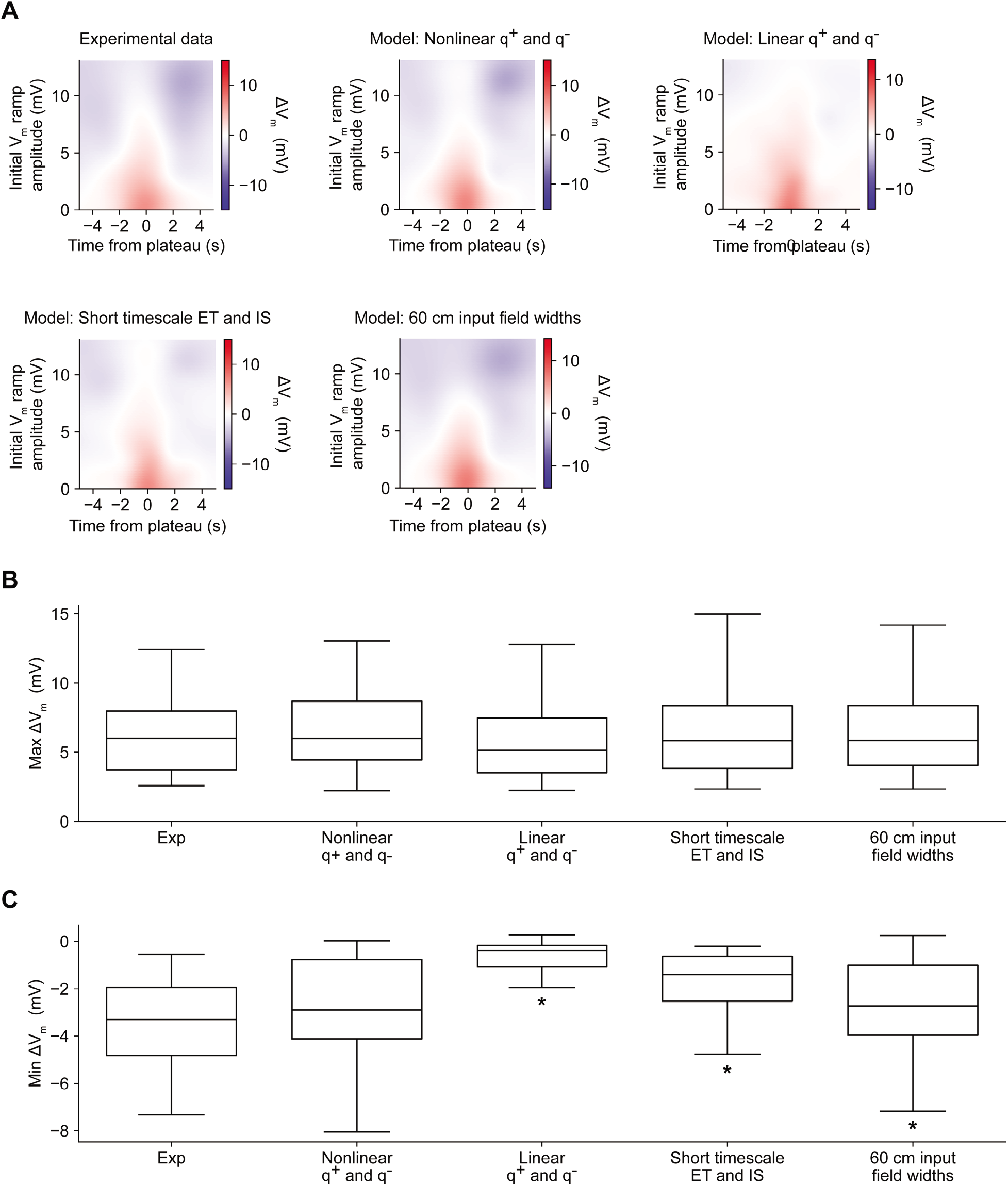
Comparison of alternative models of BTSP (related to Figures 5 and 6). (**A**) Variants of the model of bidirectional BTSP described in Figures 5 and 6 were evaluated based on the accuracy of their predictions of experimentally measured changes in *V*_m_ ramp amplitude induced by dendritic plateau potentials. For each model variant, plots show change in model *V*_m_ ramp as a function of both time and initial *V*_m_ ramp amplitude (see Materials and Methods). (**B**) The maximum change in *V*_m_ for each induction in the experimental dataset (Figures 1–3, n=26 inductions) predicted by each model is compared to the experimental data. Differences between groups were first statistically evaluated with a Friedman test (p = 0.08926). No differences were detected, suggesting that all model variants accurately accounted for the potentiation component of BTSP. (**C**) The minimum change in *V*_m_ for each induction in the experimental dataset (Figures 1–3, n=26 inductions) predicted by each model is compared to the experimental data. Differences between groups were first statistically evaluated with a Friedman test (p < 0.00001), and then model variants were compared to the experimental data with posthoc Wilcoxon signed rank tests. Asterisk indicates p < 0.05, after Bonferonni correction for multiple comparisons. See Materials and Methods for formulation details of each model.

**Figure S8.**
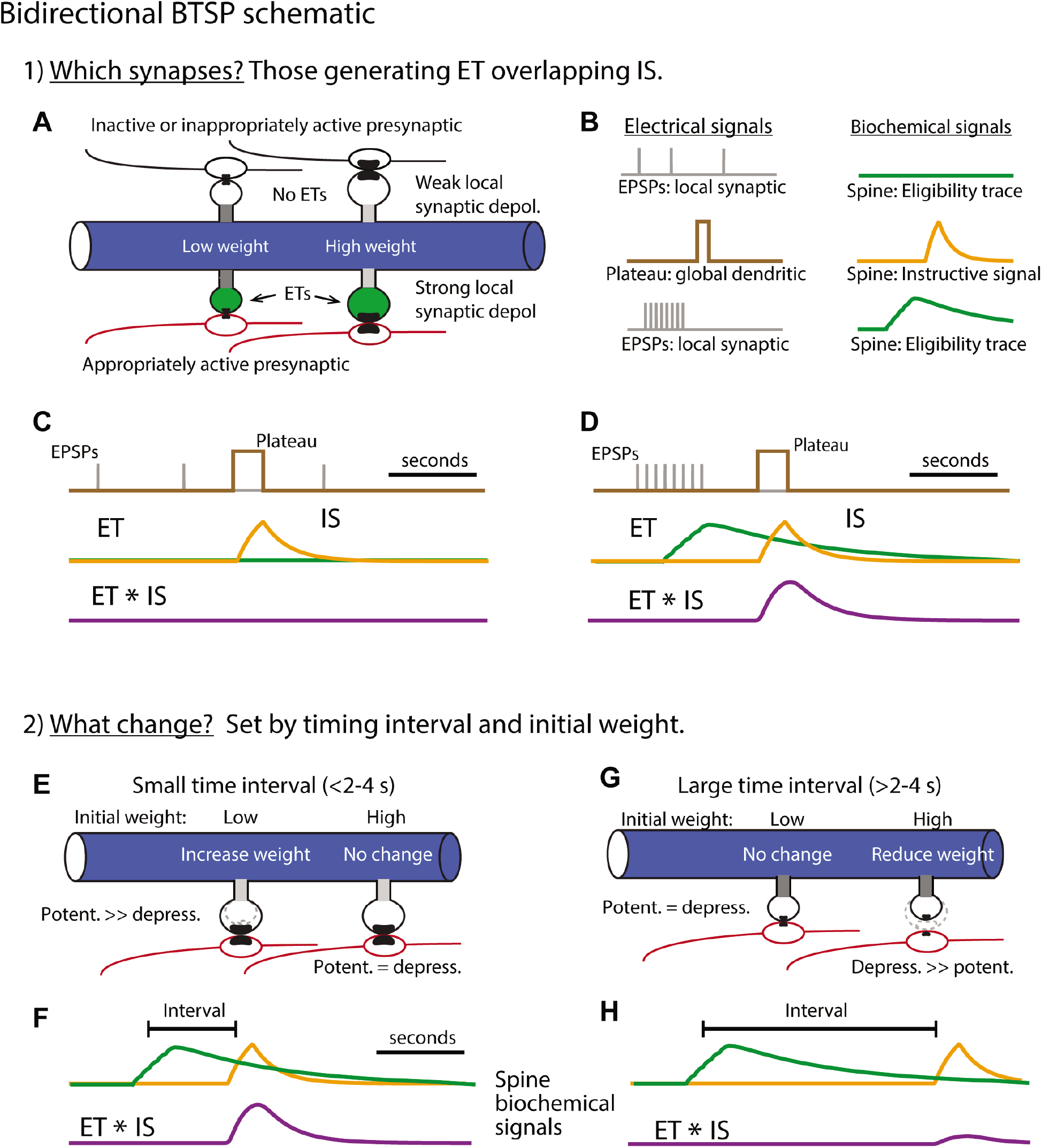
Bidirectional BTSP schematic (related to Figures 3, 5, and 6). (**A** – **C**) Step 1) Selecting synapses for weight adjustment. (**A**) Postsynaptic dendrite with spines and presynaptic inputs. Top spines receive inappropriately patterned or inactive inputs (black) while lower spines receive appropriately patterned inputs (red). (**B**) The electrical signals involved are i) excitatory postsynaptic potentials (EPSPs; grey) that produce local synaptic depolarization, and ii) the plateau potential (brown) that produces global dendritic depolarization. Each of these two electrical signals produce a biochemical signal that prolongs their impact with EPSPs generating eligibility traces (ETs, green) and the plateau generating an instructive signal (IS, yellow). Only appropriately patterned inputs generate ETs (lower set of traces in (B) and spines shaded green in (A)). This sensitivity of ET generation to presynaptic activity pattern (i.e. input repetition, synchrony and spatial clustering) may be a result of the voltage-dependence of NMDA-R activation. (**C**) Overlay of time courses of electrical (grey and brown) and biochemical signals (green and yellow) generated by inappropriate or inactive inputs. The lack of ET generation produces no overlap signal (ET * IS, purple). Synapses lacking significant overlapping ET and IS will not have their weights adjusted. (**D**) Same as (C) but for appropriately patterned inputs. Synapses that have overlapping ET and IS (ET * IS, purple) are selected to have their weights adjusted. (**E** – **H**) Step 2) Determining the amplitude and direction of synaptic weight changes. Only spines selected in step 1 are shown. (**E** – **F**) Short intervals between synaptic inputs and plateau potentials (E) produce large magnitude overlap signals (F). Resulting weight changes are dominated by potentiation due to subsequent nonlinear potentiation and depression processes which are scaled by initial weights (see Figure 5 and Materials and Methods). An interplay between the initial input weight and the time interval determines the exact magnitude of the change in weight. (**G** – **H**) Longer intervals (G) produce smaller overlap signals (H) and depression dominates, again due to the subsequent nonlinear processes (see Figure 5 and Materials and Methods). The exact magnitude of synaptic depression is determined by an interplay between the interval and the initial weights.

**Figure S9.**
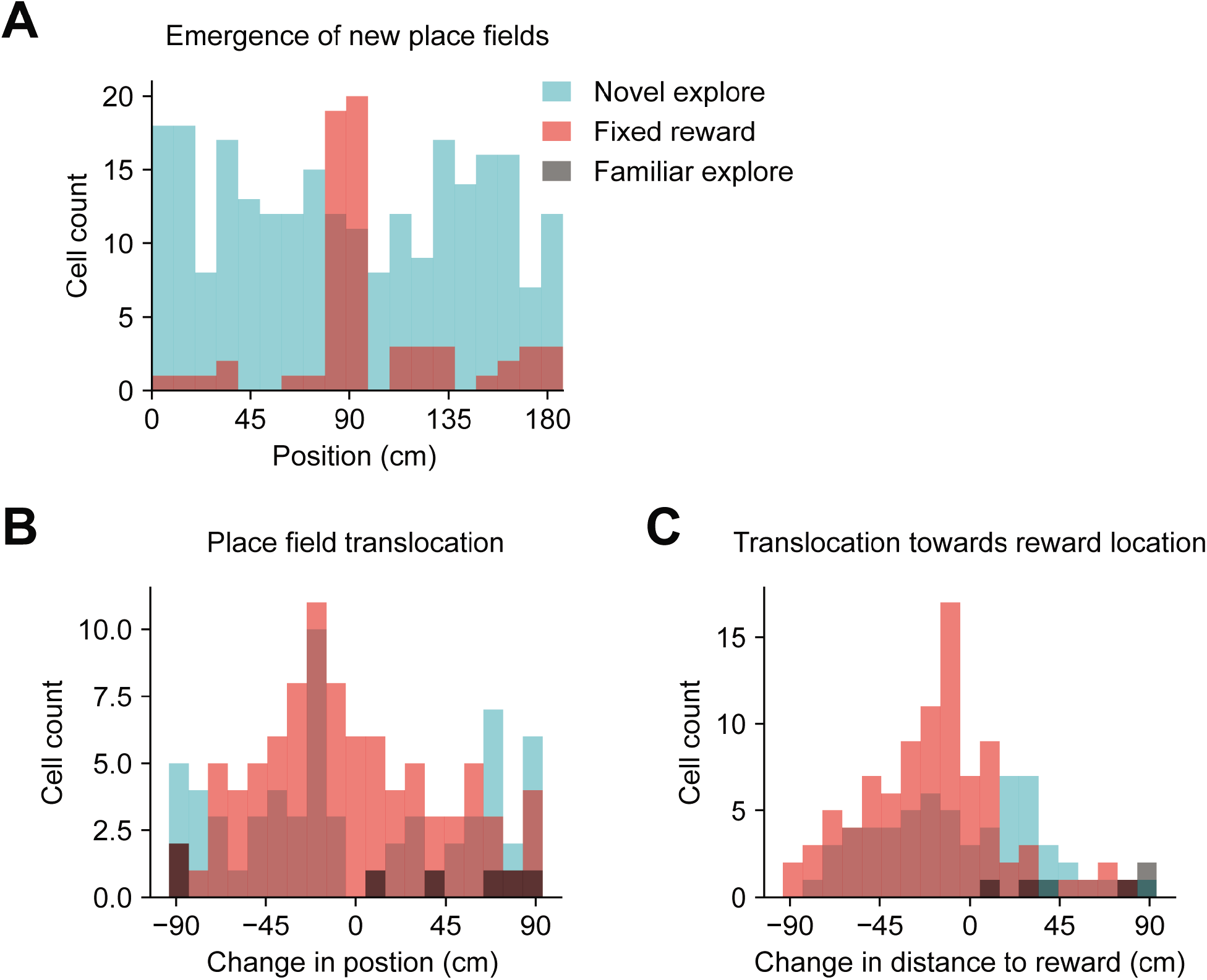
New place field acquisition and pre-existing place field translocation in a network model of goal-directed navigation (related to Figure 7). (**A** – **C**) Data shown are derived from the network model results from Figure 7. (**A**) Histogram depicts neurons recruited to express new place fields in each spatial bin (epochs: novel explore, blue; fixed goal, red; familiar explore, grey). (**B**) Histogram depicts absolute change in position of translocated place fields. (**C**) Histogram depicts change in position of translocated place fields relative to the goal location.

## Notes

### Competing Interest Statement

The authors have declared no competing interest.

### Summary of Updates

Additional analysis and modeling work now included. Results and discussion revised for clarity.

